# Integrative chromatin domain annotation through graph embedding of Hi-C data

**DOI:** 10.1101/2022.04.14.488414

**Authors:** Neda Shokraneh, Mariam Arab, Maxwell Libbrecht

**Affiliations:** Computing Science Department, Simon Fraser University, Burnaby, V5A 1S6, Canada

## Abstract

**Motivation:** The organization of the genome into domains plays a central role in gene expression and other cellular activities. Researchers identify genomic domains mainly through two views: 1D functional assays such as ChIP-seq, and chromatin conformation assays such as Hi-C. Fully understanding domains requires integrative modeling that combines these two views. However, the predominant form of integrative modeling uses segmentation and genome annotation (SAGA) along with the rigid assumption that loci in contact are more likely to share the same domain type, which is not necessarily true for epigenomic domain types and genome-wide chromatin interactions.

**Results:** Here, we present an integrative approach that annotates domains using both 1D functional genomic signals and Hi-C measurements of genome-wide 3D interactions without the use of a pairwise prior. We do so by using a graph embedding to learn structural features corresponding to each genomic region, then inputting learned structural features along with functional genomic signals to a SAGA algorithm. We show that our domain types recapitulate well-known subcompartments with an additional granularity that distinguishes a combination of the spatial and functional states of the genomic regions. In particular, we identified a division of the previously-identified A2 subcompartment such that the divided domain types have significantly varying expression levels.

**Availability:** https://github.com/nedashokraneh/IChDA

**Supplementary information:** 

## 1 Introduction

Chromosomes are segmented into different domain types that are identified through analysis of genomic data Bickmore and van Steensel (2013); Rao *et al*. (2014); Shin *et al*. (2016); Lee and Roy (2020); Filion *et al*. (2010); Sexton and Cavalli (2015); Dixon *et al*. (2016). Genome domain organization plays a central role in many cellular activities including gene expression and DNA replication Rao *et al*. (2014). The disruption of this regulation leads to developmental defects and diseases like cancer. Therefore, identifying genomic domain types provides an insight into domain-scale mechanisms behind cell differentiation and disease development.

Heitz Heitz (1928) first classified genomic regions into heterochromatin or euchromatin based on differential compaction at interphase from microscopy observations. Heterochromatin is a condensed region that is not accessible by regulatory elements, while euchromatin is less condensed, more accessible, and highly transcribed Grewal and Jia (2007).

Later, the development of functional assays enabled genome-wide mapping of different proteins and histone modifications Bickmore and van Steensel (2013). Here, we use the term “functional assay” to mean any experiment that aims to measure biochemical function or activity (such as ChIP-seq, DNase-seq, Repli-seq, and others), whether or not these marks necessarily confer biological function. These functional assays are usually represented as a 1D vector over the genome. Clustering genomic regions according to functional genomic signals at a coarse resolution (10 – 100 kb) has been shown to identify genomic domain types such as active and repressed functional domains Thurman *et al*. (2007); Larson and Yuan (2010); Filion *et al*. (2010).

Recently, chromatin conformation capture assays such as Hi-C provided a way to measure the genome-wide contact map among genomic regions Lieberman-Aiden *et al*. (2009). Analysis of Hi-C led to the identification of multiple chromatin domain structures such as compartments, TADs and loops Lieberman-Aiden *et al*. (2009); Rao *et al*. (2014); Dixon *et al*. (2012); Shin *et al*. (2016); Lee and Roy (2020); Yang and Ma (2022). Compartments are chromatin domain types where each genomic region within the same compartment has a similar interaction pattern with the rest of the genome and is spatially segregated within the nucleus Lieberman-Aiden *et al*. (2009). More detailed subcompartments have been identified from Hi-C data with a high coverage that have more distinct structural and functional properties Rao *et al*. (2014). Compartments are strongly correlated with local functional genomic data including replication timing, transcriptional activity, epigenetic patterns, and higher-order nuclear structures Liu *et al*. (2016); Nuebler *et al*. (2018).

The close association between different chromosomal measurements necessitates integrative modeling that incorporates multiple data types in order to understand genome organizational patterns. Segmentation and genome annotation (SAGA) algorithms have emerged as the predominant tool for integrative modeling of multiple 1D genomic signals Libbrecht *et al*. (2021). A SAGA algorithm such as Segway or ChromHMM assigns a label to each genomic region such that regions with the same label have similar patterns of input signals. Most SAGA algorithms employ a chain-structured probabilistic model such as a hidden Markov model (HMM) structure Day *et al*. (2007); Ernst and Kellis (2012); Hoffman *et al*. (2012).

Although any SAGA algorithm based on a multivarite hidden Markov model can jointly model different types of 1D genomic signals, this model cannot jointly model 1D functional assays and pairwise Hi-C interaction data together. To remedy this problem, two previous methods were developed to perform such joint modeling. First, Segway-GBR (graphbased regularization) Libbrecht *et al*. (2015) jointly models functional genomic signals and Hi-C intra-chromosomal significant interactions through augmenting a chain-structured probabilistic model present in Segway Hoffman *et al*. (2012) with graph-based posterior regularization. This regularization uses a graph representing a Hi-C data set to encourage spatial neighbors to receive a similar label. Second, SPIN Wang *et al*. (2021) jointly models spatial genomic signals and Hi-C genome-wide significant interactions using a hidden Markov random field (HMRF) in order to identify genome-wide spatial compartmentalization patterns. In particular, they construct the Markov random field by connecting both linear and spatial neighbors, as defined by statistically-significant Hi-C interactions. They learn emission and transition probabilities based on observed spatial signals and their spatial neighbors. This approach encourages positions which are in contact in 3D to have a similar pattern of domain labels to positions that are neighbors in 1D.

The Segway-GBR makes the core assumption that positions connected in 3D are more likely to share the same label. This assumption is motivated by the observation that domain labels are correlated in 3D, but it may not hold in all cases. The SPIN approach, HMRF, is more flexible and can learn the transition probabilities between labels instead of assuming higher transition probabilities between the same labels. However, this approach still assumes that the transition probability of connected positions is invariant genome-wide. Furthermore, in practice, the HMRF usually assigns a higher probability to connected positions sharing the same label and thus enforces the pairwise prior in the annotation process. Thus, we need a domain modeling strategy that avoids this pairwise prior assumption.

The recent development of Hi-C embedding methods provides a way to summarize pairwise chromatin conformation into a small number of 1D genomic tracks Xiong and Ma (2019); Ashoor *et al*. (2020); Dsouza *et al*. (2021). These embedding representations have been shown to be effective for many tasks including identifying subcompartments, improving Hi-C resolution and simulating genomic perturbations.

Here, we propose to leverage Hi-C embeddings to perform integrative identification of chromatin domains. Following SCI Ashoor *et al*. (2020) we use the graph embedding model LINE Tang *et al*. (2015) to embed Hi-C data into 8 1D structural features that capture genomewide compartmentalization pattern of genomic bins. We then infer combinatorial domain types by inputting functional and structural genomic signals into a hidden Markov model-based SAGA algorithm. We show that, because functional and structural compartmentalization are different genome organizational layers, this approach outperforms existing integrative approaches such as Segway-GBR and SPIN in capturing both the shared and data-specific patterns of functional and structural genomic signals. We use this approach to identify 8 combinatorial domain types and show that they recapitulate chromatin epigenomic and compartmentalization patterns simultaneously and grant additional granularity to previously-identified subcompartments.

## 2 Materials and Methods

### 2.1 Dataset

We used 12 functional assays, including ChIP-seq targeting H2A.Z and 10 histone modifications (H3K4me1, H3K4me2, H3K4me3, H3k9ac, H3K9me3, H3K27ac, H3K27me3, H3K36me3, H3K79me2, H4K20me1), and DNase-seq, as 1D functional genomic signals. We used Hi-C data to learn genomic structural features, and add spatial constraints to existing integrative models (graph construction from Hi-C data is explained in the supplementary information section 2). We also used gene expression, replication timing, and ChIA-PET data to evaluate domain annotations. All data are on the GM12878 and K562 cell lines because these cell lines have the most available data. The details of data sources and processing are provided in the supplementary information section 1.

### 2.2 Learning genome-wide global structural features

We use a node embedding model, LINE Tang *et al*. (2015), similar to the subcompartment caller SCI Ashoor *et al*. (2020) to learn genomewide global structural features representing the spatial organization of the genome. In this section, we describe the general formalization of node embedding models followed by details of the used model, LINE with second-order proximity objective.

In general, a node embedding model maps each node in the input network graph into a low-dimensional vector space (also known as an embedding space), such that pairwise distances in the embedding space are representative of the proximity of pairs of nodes in the input graph. They have four main components, (1) encoder: to map the graph structure into a vector space, (2) proximity definition: to define a proximity measure between pairs of nodes based on the input graph structure, (3) decoder: to map the embeddings of a pair of nodes to a reconstructed proximity measure, and (4) loss function: to penalize the difference between the reconstructed and true proximity measures.

The most important component to define is the proximity measure which is based on the graph structure that we aim to embed. Given a weighted graph *G* = (*V, E*), where *V* is the set of nodes (*v_i_* , …, *v*_*n*_), and *E* is the set of edges attributed with a *w_ij_* representing the weight of the edge between nodes *v_i_* and *v_j_* , the first-order proximity of a pair (*i, j*) is equal to the weight *w_ij_* of the edge connecting that pair. A greater edge weight between nodes indicates more first-order proximity between nodes.

A greater similarity between nodes’ interaction patterns with the rest of the nodes in a graph indicates a higher second-order proximity between the two nodes. More formally, a first-order proximity of a node *v_i_* with all other nodes is defined as *p_1_*(*v_i_*) = (*w*_*i*1_, …, *w_in_*), and the second-order proximity of a pair (*i, j*) is defined as the similarity between *p*_1_(*v_i_*) and *p*_1_(*v_j_*).

LINE can be optimized based on either first-order or second-order proximity loss functions. Since we aim to capture the global chromatin structural features, we will use LINE embeddings based on the second-order proximity. We also show that second-order embeddings capture more meaningful global chromatin structural patterns compared to first-order embeddings in the supplementary information section 3. Here, we describe the LINE model based on the second-order proximity in more detail.

To learn the embeddings that capture the second-order proximity between nodes, two vectors corresponding with each node *v_i_* are defined, the embedding of the node, *u_i_*, and the context of the node, 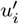, that comes from the embedding of its neighbors. Then, *p*_2_(.|*v_i_*) is defined for each node *v_i_* as a distribution over all the nodes that determine the probability that the context of each node has been generated using embedding of node *v_i_*.

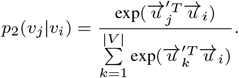

To preserve the second-order proximity, *p*_2_(.|*v_i_*) should be close to the empirical distribution coming from the graph structure, defined as 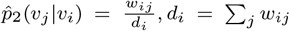. The objective function aims to minimize the weighted sum of distances between *p*_2_(.|*v_i_*) and 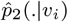 over all the nodes, defined according to the Kullback-Leibler divergence between *p*_2_(.|*v_i_*) and 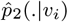. A weight λ_*i*_ is given to each node *i* equal to its degree. The full objective function is defined as

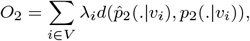

which is simplified to:

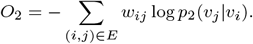

We use LINE’s Tang *et al*. (2015) original implementation to learn the global chromatin structural features from O/E chromatin interaction graph (supplementary information section 2). The optimization strategies used in their implementation makes it applicable to large graphs like genome-wide chromatin interaction graph. Details about choosing the node embedding model and hyperparameters selection are provided in the supplementary information section 3.

### 2.3 Domain annotation using segmentation and genome annotation (SAGA)

Our approach involves applying existing segmentation and genome annotation (SAGA) methods to integrate 1D and 3D genomic features. Unsupervised segmentation and genome annotation (SAGA) algorithms cluster the genomic positions based on their observed functional or chemical genomic signals Libbrecht *et al*. (2021) and assign a label to each cluster based on biological interpretation of genomic signals pattern. These algorithms have most commonly been applied at a fine resolution to define 100 – 1000 bp segments, resulting in annotations of smallscale phenomena such as protein-coding genes and regulatory elements. However, these algorithms can also be applied on genomic signals defined on larger segments (10^4^ – 10^6^ bp) to find domain-scale patterns and labels such as Polycomb-repressed domains, heterochromatic domains and different types of active domains Filion *et al*. (2010); Bickmore and van Steensel (2013); Libbrecht *et al*. (2015); Marco *et al*. (2017).

SAGA algorithms use structured probabilistic models that model the dependency between latent labels and observed genomic signals, and the dependency between neighbor latent labels on a genome sequence. We use following methods as a baseline: (1) Gaussian Mixture model (GMM), the most simple SAGA method that only models the dependency between latent labels and observed genomic signals, (2) Hidden Markov model with observations modeled as GMM (GMM-HMM) that model the dependency between neighbor latent labels, (3) Segway Hoffman *et al*. (2012), which is specifically designed for modelling genomic signals, and (4) Segway-GBR Libbrecht *et al*. (2015) and (5)SPIN Wang *et al*. (2021), which extend existing SAGA algorithms to incorporate the dependency between spatial neighbors as well (more details are provided in the supplementary information section 6).

### 2.4 Combinatorial HMM: an integrative approach to model 1D signals and spatial dependencies

By applying a node embedding model to genome-wide Hi-C data, we map each genomic bin into a vector space. A vector space with dimension 8 is enough to capture the structural information in whole Hi-C data (supplementary information section 3).

Then, we input the learned 8 structural features with 12 functional genomic signals to HMM, which we showed that it is a proper choice for the domain-scale annotation (section 3.1), and infer domain types that we call HMM_combined. We also input the learned structural features and functional genomic signals to HMM separately to identify HMM_structural and HMM_functional domain types separately as baselines.

### 2.5 Evaluation methods

#### 2.5.1 The Percentage of Variance Explained metric

The percentage of variance explained (VE) is used to measure the consistency between different epigenomic signals, such as histone modifications and gene expressions, with annotated domains. The VE is a fraction of signal variance that is explainable using an annotation prediction. Given a genome annotation *a*_1:*n*_ ∈ {1…*K*}^*n*^ (*K* is the number of labels) and a signal vector *s*_1:*n*_ ∈ ℝ^*n*^, the mean value of the signal over positions with label *k* is calculated as

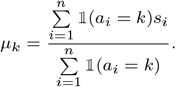

So, the predicted signal vector is 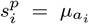. VE is computed as the difference of total variance var(*s_i_*) and the variance of residuals of the prediction var(*d_i_*), where 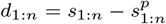

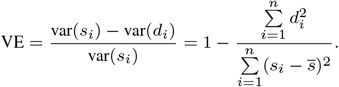

VE is bounded by the range [0, 1], and higher values indicate more agreement between annotation and signal values. The bootstrap VEs to estimate their standard errors is explained in the supplementary-information section 5.1.

#### 2.5.2 Gene expression

The most important function of domain structures is transcription regulation, and we expect that domain states have different transcriptional patterns. We acquired RNA-seq measurements of gene expression, measured in RPKM (supplementary information section 1). We annotate each gene with the most frequent domain label among the gene region and calculate the proportion of variance explained of gene expression values given domain labels as gene expression VE. To reduce the influence of large outlier gene expression values, we transform gene expression values using the inverse hyperbolic sine function 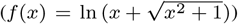 before calculating a VE.

#### 2.5.3 Replication timing

The order with which DNA is replicated in S phase is correlated with many genomic phenomena including domain regulation Rhind and Gilbert (2013). Thus, we used 6-phase replication timing (RT) data to evaluate the domain annotations. The RT data is in 1000 bp resolution, such that each 1000 bp-length region is replicated with a proportion *p_i_* in a phase *i*.

First, we change the resolution of RT signals from 1 kb to 100 kb resolution to match the annotation’s resolution. Then we calculate the VE for each phase separately and calculated a mean of VE among 6 phases as RT VE.

#### 2.5.4 Agreement with ChlA-PET loops

We also assessed the domains with respect to higher-order chromatin structures, including loops from CTCF and RNAPII Heidari *et al*. (2014) ChIA-PET data. If domain structures capture the higher-order structure of the chromatin, we expect ChIA-PET loops to usually have the same label on their ends. Therefore, we grouped loops based on the labels assigned to their ends, resulting in a matrix, *O_L×L_*, where *L* is a number of labels, *O_i_j* is a number of loops with one end belonging to the label *i*, and the other belonging to the label *j*. Given a normalized coverage of labels *C* = (*C*_1_, …,*C_L_*), the expected number of loops between labels *i* and *j* is *E_ij_* = total_*O*_ * *C_i_* * *C_j_* , where total_*O*_ = Σ*_i,j_ O_ij_* . The expected and observed number of loops with a same label on their ends are Σ*_i_ E_ii_* and Σ*_i_ O_ii_*, respectively. The ratio of observed over expected number of loops with a same label shows the enrichment of loops occurring between similar domain states. The higher ratio indicates the higher consistency between the domain annotation and the higher-order structure input. We call observed over expected enrichment of CTCF and RNAPII ChIA-PET loops as CTCF OE and RNAPII OE scores respectively. The bootstrap OE scores to estimate their standard errors is explained in the supplementary section 5.2.

### 3 Results

#### 3.1 Functional and structural features confer complementary information

To ascertain the need for combinatorial domain annotations, we evaluated whether features of functional and structural domain types confer complementary or equivalent information. To do so, we created domain annotations for cell types GM12878 and K562 using each of functional and structural chromatin features, and we found that they confer complementary information. In particular, we found that structural domains are broader than functional domains and they regulate chromatin functions in domain-scale.

To identify domains based on functional features, we applied three baseline annotation models (GMM, HMM, and Segway) on functional genomic signals, and identify 6 domain types for GM12878. The average length of Segway domains (0.4 Mb) is longer than HMM domains (0.25 Mb) and HMM domains are also longer than GMM domains (0.2 Mb) due to the HMM and Segway duration models (Fig. S5a). We compared these three models in terms of explaining different cellular activities and chromatin features described in the evaluation methods (Fig. S5a) and found that HMM domains are the best in 3/4 scores, therefore, we chose HMM as an annotation approach for identification of different domain types. We also chose 6 domain types based on the proportion of variance explained (VE) for input signals (Fig. S5b), as VE does not improve significantly for any of the input signals for the number of domain types greater than six. This is also consistent with known chromatin types in a domain-scale Bickmore and van Steensel (2013); Filion *et al*. (2010); Libbrecht *et al*. (2015).

To identify domains based on structural features, we learned genomic structural signals using the LINE embedding model for all genomic bins in a shared space (section 2.2), applied an HMM on the learned structural signals, and identified 6 domain types (HMM_structural).

We named HMM_functional and HMM_structural domain types based on their overlap with subcompartments from HF1 to HF6 and HS1 to HS6, respectively, such that the overlap with inactive subcompartments increases by increasing the domain type’s number (Fig. S6 and Fig. S7 for GM12878 and K562 cell types, respectively). We use the fold-change of each domain type from the first annotation over each domain type from the second annotation to calculate the overlap between domain types from the first and second annotations (supplementary information section 4).

We compared HMM_functional, HMM_structural, and subcompartment annotations for GM12878 cell type in terms of their domain sizes, the overlap of their domain types, and explanation of other chromatin features including gene expression and replication timing profiles and loops from ChIA-PET data (Fig. 1). We observed that structural domains are broader than functional domains (Fig. 1a), and there is less number of structural domains. The structural domain types are highly similar to subcompartments (Fig. 1b), which confirms that the learned structural signals capture a genome-wide structural compartmentalization pattern. B2 and B3 subcompartments are only mixed and divided in another way into HS4, HS5, and HS6. This is also observable from structural embeddings UMAPs that B2 and B3 subcompartments are not distinguishable by learned structural embeddings (Fig. S2a). On the other hand, functional domain types are less similar to subcompartments and HMM_structural domain types. HF1 and HF2 overlap with both active subcompartments (Fig. 1b), and the other four HMM_functional domain types are mixes of A2 and other inactive subcompartments.

**Fig. 1:**
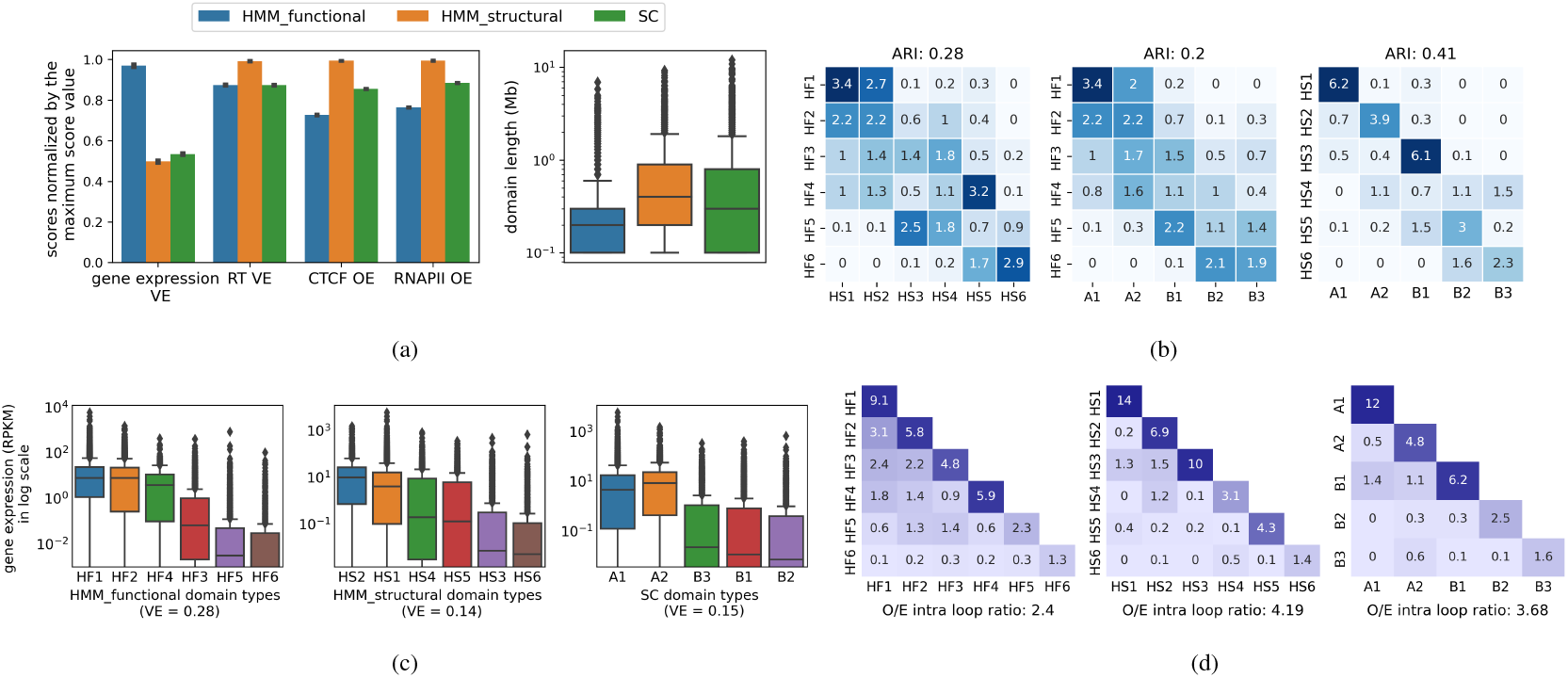
(a) Comparison of HMM_functional, HMM_structural, and subcompartments Rao *et al*. (2014) domain types based on the proportion of variance explained for gene expression (gene expression VE) and replication timing signals (RT VE), agreement with CTCF and RNAPII ChIA-PET loops (CTCF OE, RNAPII OE), and the distribution of domain lengths. (b) The overlap between each pair of HMM_functional, HMM_structural, and subcompartments domain types and similarity (adjusted rand index (ARI)) between the annotations. (c) The distribution of gene expression values in RPKM for each domain type of HMM_functional, HMM_structural, and subcompartments annotations. ’VE’ in titles means the variance explained for gene expression given a domain annotation. (d) The observed/expected CTCF ChIA-PET loops between each pair of labels for different annotations.

The chromatin features that are regulated in domain-scale such as replication timing and ChIA-PET loops are better explained by structural domain types (Fig. 1a). We calculated the enrichment of chromatin loops between every two domain types given a domain annotation, which is calculated as the ratio of observed loops over expected counts between two domain types, such that the expected count is relative to the coverage of two domain types. Most of the loops are between pairs with the same domain type given HMM_structural and subcompartments annotations, while there are more loops between different domain types in HMM_functional annotation (Fig. 1d). We summarized the enrichment of loops with ends from the same domain type into the observed/expected (OE) score, which estimates the agreement of loops with annotation. We expect the OE score to be high, and the ratio of HMM_structural to HMM_functional OE scores for CTCF and RNAPII loops are 1.36 and 1.3 respectively (Fig. 1a).

In contrast, a more local feature, gene expression, is better explained by HMM_functional annotation (Fig. 1a). There is more variation of gene expression values between domain types compared to within domain type given HMM_functional annotation, while there is less between-domain type variation between HMM_structural and subcompartments domain types (Fig. 1c). For example, the two-sample t-test between gene expression of two active HMM_functional (HF1 and HF2), HMM_structural (HS2 and HS1), and subcompartments (A1 and A2) domain types result in p-values 2.87e–6, 0.035 and 0.97, respectively.

Due to the availability of structural genomic signals, LaminA and SON TSA-seq Chen *et al*. (2018), for the K562 cell type, we also compared different domain types based on the fold enrichment of structural genomic signals (Fig. S7). The proportion of variance explained for LaminA and SON TSA-seq signals given HMM_structural annotation are 91% and 92% higher than HMM_functional annotation (Fig. S8).

These observations show that the separation of active functional domain types is based on their local enrichment of functional activities such as histone modifications, while the separation of active structural domain types is mostly based on their positioning within the nucleus as measured indirectly by Hi-C. Therefore, functional and structural compartmentalization of chromatin are different chromatin organization layers and complementary to each other, and thus an integrative approach is needed to model both patterns together.

#### 3.2 Combinatorial domain types better capture the chromatin higher-order organization compared to existing integrative models

To infer domain types that jointly capture chromatin functional and structural compartmentalization, we input functional and learned structural genomic signals to the HMM (section 2.4); we term the resulting combinatorial annotation HMM_combined.

We compared the HMM_combined annotation to existing methods that utilize both measurements of 1D genomic signals and 2D Hi-C data, Segway-GBR and SPIN. The graph-based posterior regularization (GBR) can be added to any type of probabilistic model, and we use the GMM_GBR version using GMM as a base probabilistic model to be consistent with two alternatives, SPIN, and HMM_combined, that utilize GMM as their basic component as well. Note that by SPIN annotations, we mean the result of applying HMRF, the approach used for SPIN, to our datasets.

We ran HMRF, SPIN’s method, on our data, functional genomic signals, and significant Hi-C interactions from GM12878 cell type. We observed that SPIN results in just two common labels with coverage of 0.5 and 0.49 (Fig. S9b), even though SPIN was run with *k* = 6. Other labels have very low coverage and are not interpretable, therefore, we made SPIN annotation with two labels HMRF1 and HMRF2 (to be not confused with original SPIN states), where HMRF1 is active and HMRF2 is inactive. Further analysis of SPIN result is provided in the supplementary information section 7. The GMM_GBR domains have a high agreement with HMM_functional domain types (Fig. S11c), and do not have a higher agreement with subcompartments and compartments compared to the original HMM_functional domain types (Fig. S12a). We also named GMM_GBR annotation labels from GBR1 to GBR6 based on their enrichment of active to inactive subcompartments. HMM_combined domains have a high agreement with both HMM_functional domains, and subcompartments (Fig. S12a), showing that it captures both local chromatin activities and higher-order structure simultaneously. Six HMM_combined domain types are named HC1 to HC6 based on their enrichment of active to inactive subcompartments. Detailed characterization of HMM_combined, GMM_GBR, and SPIN annotations labels and their overlap with other annotations are shown in Fig. S10 and Fig. S11.

Then, we compared three integrative models in terms of explaining different cellular activities and chromatin features (Fig. 2a). We can see that HMM_combined outperforms both SPIN and GMM_GBR in all metrics except for the proportion of variance explained for the gene expression. This is due to the high similarity of the GMM_GBR annotation to the HMM_functional annotation. Thus, GMM_GBR mostly captures local chromatin types that are sufficient to explain the local cellular activities, such as gene expression. We found similar results in the K562 cell type as well (Fig. S14), except that in this cell type we did not get a valid annotation from SPIN (one label covered 97% of the genome) and therefore we excluded SPIN from the analysis. This limitation is likely due to the strong pairwise prior assumption made by GMM_GBR, which assumes that positions close in 3D are more likely to be of the same domain type, and lack of clear pattern between the labels of spatial neighbors, such that in our experiments, SPIN’s HMRF model produced poor results (supplementary information section 7). Therefore, it is necessary to embed the higher-order organization of the chromatin into a vector space at first, to preserve the information of both chromatin local types and higher-order conformation in inferred domain types.

**Fig. 2:**
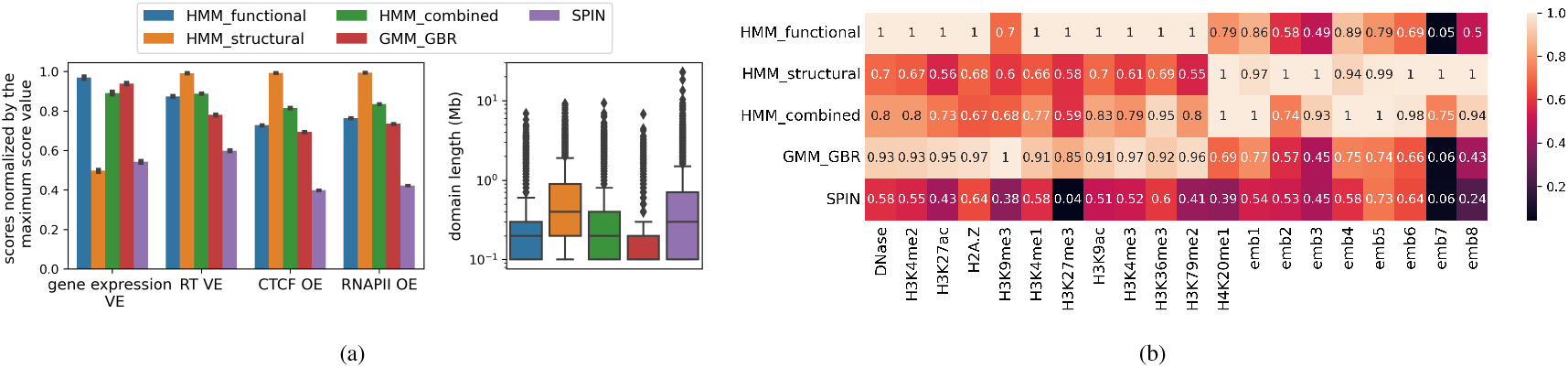
(a) Comparison of HMM_functional, HMM_structural, HMM_combined, GMM_GBR, and SPIN annotations based on the proportion of variance explained for gene expression (gene expression VE), the average proportion of variance explained for replication timing 6 phases signals (RT VE), agreement with CTCF and RNAPII ChIA-PET loops (CTCF OE, RNAPII OE), and the distribution of domain lengths. (b) The normalized proportion of variance explained for 12 functional genomic signals and 8 structural embeddings given different annotations, where VEs for each signal are normalized by the maximum VE among different annotations.

To find the contribution of different genomic signals in different annotations, we calculated the normalized variance explained for 12 functional genomic signals and 8 structural embeddings given different annotations (Fig. 2b) for GM12878 cell type. The HMM_combined annotation captures higher-order conformation with a significantly better explanation of functional genomic signals compared to HMM_structural annotation. Furthermore, it is observable that structural features have more contribution to HMM_combined annotation, which is also confirmed by the higher similarity between HMM_combined and HMM_structural annotations compared to between HMM_combined and HMM_functional annotations (Fig. S12a). However, this observation is not generalizable to all cell types (Fig. S12b). This might be the result of the quality of learned structural features from the K562 cell type due to its lower read depth (Fig. S13).

#### 3.3 Combinatorial domains capture different chromatin organization layers

We found that the domain types learned by the combinatorial HMM recapitulate subcompartments and identify additional granularity among these compartment types. As a result of the aggregation of functional and structural genomic signals, it is expected to find more than 6 domain types. Therefore, we evaluated combinatorial domain annotations with 6-10 types according to gene expression VE and RT VE (Fig. S8). We choose 8 domain types because gene expression VE does not change by increasing the number of domain types, RT VE’s change is small, and we were not able to interpret further domain types, as discussed later. We named HMM_combined domain types from HC1 to HC8 based on their enrichment of functional signals and subcompartments, and to further our understanding of the inferred combinatorial domain types, we compared them to functional genomic signals, overlap with subcompartments, average gene expression, number of genes per 1 Mb region, and their coverage along the genome (Fig. 3).

**Fig. 3:**
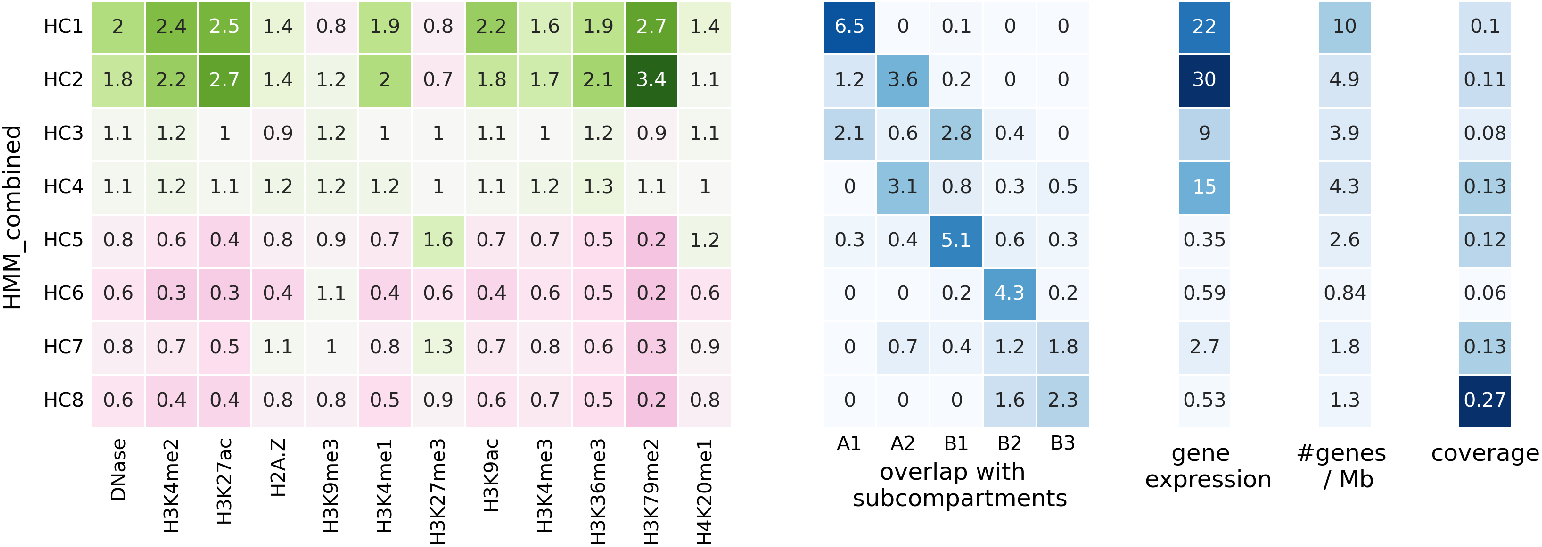
The fold enrichment of genomic signals (column 1) and subcompartments (column 2), an average of transcription level of genes (column 3), the density of genes (column 4) and coverage (column 5) of each of 8 domain types from HMM_combined annotation.

These eight combinatorial domain types can be classified into 4 active (HC1-HC4) and 4 inactive types (HC5-HC8). These types largely recapitulate known subcompartments. HC1 corresponds to the A1 subcompartment (associated with broad transcription) and is enriched with transcribed genes.

Combinatorial domains reveal a distinction between two types of A2 subcompartments (associated with cell type-specific transcription), represented by HC2 and HC3. HC2 overlaps with A2 and has a lower gene density compared to HC1; however, its genes are highly transcribed. HC3 also overlaps with A2, however, it is less enriched with active marks, and its genes are moderately expressed compared to HC2. It shows that the A2 subcompartment can be further divided into two domain types with different epigenomic and gene expression patterns.

HC4 is the last active type, which has a lower coverage and is less transcribed compared to other active types. These domains are mixed between the A1 and B1 subcompartments and feature moderate activating signals and gene expression. This domain type confirms the existence of genomic regions interacting with the active compartment but without high transcription.

The other four combinatorial labels correspond to inactive domain types. HC5 corresponds to the B1 subcompartment (associated with facultative repression), and it is highly enriched with H3K27me3, representative of facultative heterochromatin Bickmore and van Steensel (2013); Libbrecht *et al*. (2015). It includes a moderate number of genes; however, they are highly inactivated.

HC6 corresponds to the B2 subcompartment (associated with nucleolar domains): it is enriched with the H3K9me3 mark, representative of constitutive heterochromatin, and depleted from genes.

Combinatorial domains also reveal a distinction between two types of constitutively inactive subcompartments (B2 and B3) (associated with constitutive repression). HC7 is enriched with both inactive marks, H3K27me3 and H3K9me3, while HC8 is depleted of all epigenomic marks.

We also visualized the same plot for HMM_functional and HMM_structural 8 domain types (Fig .S15). We found that this distinction of overlap with subcompartments and gene expression pattern are not identified by HMM_functional and HMM_structural domain types respectively. These observations show that there is a link between 1D functional domain types and 3D compartment domains Bickmore and van Steensel (2013); Di Pierro *et al*. (2017), and they can supplement each other through integrative analysis.

To find the type of granularity added to combinatorial domain types through increasing the number of domain types, we identified combinatorial HMM annotations with 9 and 10 domain types. We found that the new domain type, HC9, has a low coverage of 0.03, is not enriched with a specific subcompartment (Fig. S16a), is a combination of HC3 and HC6 (Fig. S16c), and is not interpretable. Furthermore, HMM_combined with 10 domain types divide both A1 and A2 into two domain types (Fig. S16b), such that HC1 and HC2 correspond to A1, and HC3 and HC4 correspond to A2. However, the significance of the difference between gene expression patterns of HC1 and HC2 (p-value = 0.09) is much smaller than between HC3 and HC4 (p-value = 1.4e–17).

### 4 Discussion and Conclusion

The link between 1D and 3D organization of the chromatin have been studied in different scales through integrative modeling Wang *et al*. (2021); Libbrecht *et al*. (2015); Belyaeva *et al*. (2017), and predictive models Fortin and Hansen (2015); Di Pierro *et al*. (2017); Huang *et al*. (2015). In this study, we focus on integrative modeling of functional genomic signals and chromatin conformation from Hi-C data to identify chromatin domain types that capture both chromatin epigenomic and conformation patterns.

Previous integrative models, Segway-GBR and SPIN, Wang *et al*. (2021); Libbrecht *et al*. (2015) relied on making assumptions about the pairwise patterns of domain types. These methods are sensitive to the procedure we use to determine spatially close genomic regions, as they will push non-colocalized regions to get the same label if we incorporate random contacts. On the other hand, methods that infer genome subcompartments are based on clustering genomic regions’ interaction patterns, which is equivalent to the second-order proximity of the nodes (genomic regions) in a Hi-C graph. Therefore, we propose a framework that has two steps, learning structural features from Hi-C data that preserve second-order proximity of genomic regions in a Hi-C graph, and using functional and learned structural genomic signals to infer chromatin combinatorial domain types. We identify 8 combinatorial domain types that recapitulate well-known subcompartments Rao *et al*. (2014) with additional granularity and have distinct patterns of different cellular activities such as gene expression and replication timing profile.

One problem with joint modeling of 1D and 3D chromatin domain organization is a gap between their spatial resolutions. For example, functional genomic signals such as ChIP-seq are available in finer resolution and their domain organization can be studied at a fine resolutions (5-100 kb), while chromatin 3D domain types, or subcompartments, have been studied in coarse resolutions (≥ 100 kb). Such a gap can be reduced by higher resolution Hi-C data or more powerful embedding models to learn embeddings in finer resolution. For example, we use both intra and inter-chromosomal Hi-C data to learn genome-wide embeddings in a shared space. The inter-chromosomal Hi-C data are very noisy in resolutions finer than 100 kb, so the learned structural features are limited to this resolution. One future work is to learn genome-wide structural features in finer resolutions through more powerful embedding approaches. Furthermore, all 12 assays that we used in our experiments are not available for all cell types. Choosing the functional assays as input and imputation of missing assays is another future work. Finally, hierarchical hidden Markov models were proposed to identify multi-scale chromatin states including bin-scale and domain-scale from epigenomic signals Marco *et al*. (2017); Larson *et al*. (2013). Due to the multi-scale chromatin 3D organization, developing an integrative multi-scale chromatin annotation algorithm is another future direction.

## Acknowledgements

We thank the reviewers for their accurate and constructive comments.

