## Supplementary Information for "Integrative chromatin domain annotation through graph embedding of Hi-C data"

### 1 Data sources and processing

**Functional assays** We downloaded ChIP-Seq data sets targeting H2A.Z and 10 histone modifications (H3K4me1, H3K4me2, H3K4me3, H3k9ac, H3K9me3, H3K27ac, H3K27me3, H3K36me3, H3K79me2, H4K20me1), and DNase-seq data fold-change files from the Roadmap Epigenomics data portal (Table. S 1). The fold-change ratio of ChIP-Seq or DNase counts is generated using a uniform signal processing pipeline, such that the raw counts are normalized by expected background counts defined based on an Input control [1]. To calculate a signal value for each genomic bin, we calculate the average signal value over that bin. We also considered two alternative signal representations: p-value signal and the number of broad-domain peaks. We binned these tracks in the same way. Fig. S 1 shows that all three processed signals are correlated, however, the number of peaks is discrete and less informative. We choose fold-change because the average of fold-change values is more reasonable than p-values, and it is the choice of the well-known existing annotation method, Segway [2]. Note that each fold-change, p-value, and peak count data for ChIP-Seq assays are available in the mentioned link in Table. S 1.

**ChIA-PET data** We used existing processed contact lists using ChIA-PIPE available in $4DNES7IB5LY9$ [3] and $GSE59395$ [4](Table. S 1) for GM12878 and K562 cell types respectively. The processed files from $4DNES7IB5LY9$ were based on hg38 assembly, so we used lift tool to convert the genome coordinates from hg38 to hg19. After mapping $4DNES7IB5LY9$ loops to the 100 kb resolution bins, they are selected if there are more than 1 loops between bin pairs.

**Replication timing data** We used Repli-seq signals for 6 phases, G1, S1, S2, S3, S4, and G2, from Encode project (Table. S 1).

**Gene expression data** Gene expression data (in RPKM) is obtained from a Roadmap project (Table. S 1). Columns E116 and E123 correspond to gene expressions in the GM12878 and K562 cell types. Genes info such as their positions is also available in a file with the .gene_info extension in the same directory.

**Hi-C data** were obtained from [5] with GEO entry 63525. Then, observed and observed or observed over expected (O/E) contact map files for every pair of chromosomes at different resolutions were extracted using the Juicer tool [6]. The subcompartments annotation for GM12878 was also obtained from this GEO, and for K562 was obtained from SNIPER [7] GitHub repository (Table. S 1).

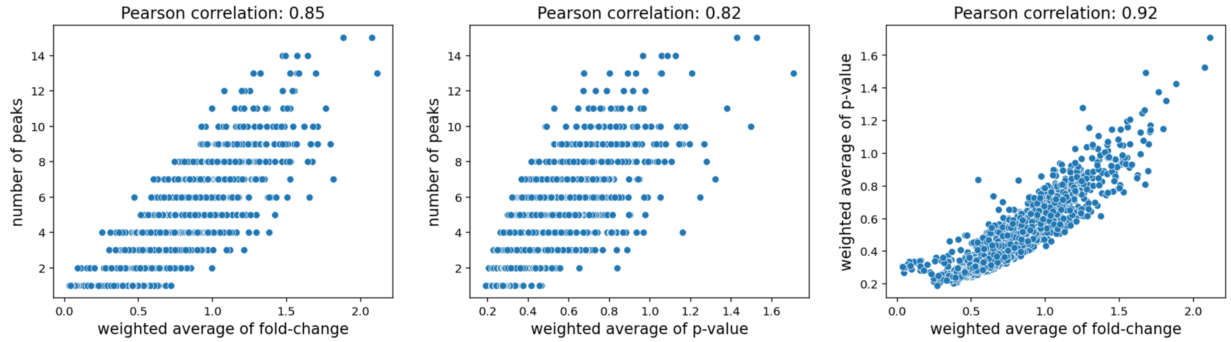


Fig. S1: Comparison of three H2A.Z ChIP-Seq processed signals: the relationship between (left) number of peaks and weighted average of fold-change, (middle) number of peaks and weighted average of p-value, and (right) weighted average of p-value and weighted average of fold-change over genomic bins (100 kb resolution) of GM12878's chromosome 1.

Table. S 1: Data sources.

| Data type | Cell type | link |
|---|---|---|
| ChIP-Seq data | GM12878 | 'E116-*.bigwig' files |
| | K562 | 'E123-*.bigwig' files |
| Hi-C data | GM12878 | hic file: 'GSE63525_GM12878_insitu_primary_30.hic' file<br>subcompartments annotation file:<br>'GSE63525_GM12878_subcompartments.bed.gz' file |
| | K562 | hic file: 'GSE63525_K562_combined_30.hic' file<br>subcompartments annotation file:<br>'K562_track.bed' file |
| ChIA-PET data | GM12878 | CTCF: 4DNFIA3H6KXY<br>RNAPII: 4DNFIDW5Q7D4 |
| | K562 | RNAPII: *.txt.gz files |
| Repli-Seq data | GM12878 | wgEncodeUwRepliSeqGm12878*bPctSignalRep1.bigWig files |
| | K562 | wgEncodeUwRepliSeqK562*PctSignalRep1.bigWig files |
| genes expression data | GM12878 & K562 | expression file: 57epigenomes.RPKM.pc.gz<br>genes info file: Ensembl_v65.Gencode_v10.ENSG.gene_info |

### 2 Chromatin interaction graph construction from Hi-C data

We need to construct the chromatin interaction graph to use as an input to (1) alternative methods, GBR [8] and SPIN [9], to represent the dependency between latent variables, and (2) a graph embedding component to capture the global chromatin structural properties.

For the first purpose, Segway-GBR uses significant Hi-C interactions called by Fit-Hi-C algorithm [10], and SPIN fit the Weibull distribution to observed over expected (O/E) Hi-C contact counts and uses significant interactions with a p-value greater than a specific threshold. While GBR annotation is less impacted by a regularization graph structure, we found that SPIN performance with ChIP-Seq data is highly dependent on the number of interactions in its Markov random field (Fig. S 9c). And, using Fit-Hi-C or Weibull p-values as a threshold results in an unbalanced number of interactions from different pairs of chromosomes that affect SPIN performance. Therefore, we use edges number as a threshold to choose significant interactions. For example, given a threshold $t$, we choose $t$ interactions with the largest O/E values (intra O/E chromatin interaction graph) or smallest Fit-Hi-C p-values (intra Fit-Hi-C chromatin interaction graph). We choose $t$ relative to the size of chromosomes for each pair of a chromosome, such that the density of edges would be the same along the genome. Note that we just use intra-chromosomal interactions for both GBR and SPIN, since both methods result in invalid and very long segments annotated with a small number of states (1 or 2) given genome-wide interactions due to the over-smoothing problem.

For the second purpose, we need to construct the interaction graph as input to graph embedding methods. This graph must eliminate local and chromosomal biases. Local biases are due to local substructures such as loops and TADs, and chromosomal biases are due to different densities of interactions in different pairs of chromosomes. To remove the local biases, we use O/E Hi-C matrices or Fit-Hi-C algorithm outputs that eliminate local biases, and for removing chromosomal biases, we extract a balanced number of significant interactions from each pair of chromosomes. We choose 2000 interactions with the largest O/E values (or smallest Fit-Hi-C p-values) from chromosome 21 (the smallest chromosome) intra-chromosomal contact map. For other pairs of chromosomes $i$ and $j$, we chose $n$ interactions, where $n = \frac{\text{size(chromosome i)} \times \text{size(chromosome j)}}{\text{size(chromosome 22)}^2} * 2000$. Then, we construct a genome-wide binary chromatin interaction graph using called significant interactions from all pairs of chromosomes, which we call the O/E (or Fit-Hi-C) chromatin interaction graph.

### 3 Model and hyperparameter selection for the graph embedding component

We choose a node embedding model, LINE [11], because it is scalable to large graphs, and has an objective based on the second-order proximity that allows capturing the global structures of the graph. Also, SCI [12] has shown that LINE performs better than alternative graph embedding models, HOPE [13] and DeepWalk [14], that capture higher-order structures of the graph in their embeddings.

Previous studies on compartmentalization used genomic bins' interaction patterns with the rest of the genome to infer their compartment labels [15, 5, 7, 12]. This is equivalent to the definition of second-order proximity (explained in the main script). We learned embeddings that preserve either first-order or second-order proximity and plotted the scatterplot of embeddings UMAPs [16] colored by subcompartment labels from Rao et al [5] (Fig. S 2a). The embeddings based on the second-order proximity are highly representative of the compartmentalization pattern as expected.

We also used the Silhouette Index (SI) [17] to assess the learned embeddings with respect to the subcompartment labels quantitatively. Given a set of points in the embedding space and their corresponding clustering labels, SI measures how similar the points are to their own cluster compared to other clusters, and ranges from $-1$ to 1, where higher values indicate that the points are clustered well. Fig. S 2b shows the comparison of SI between embeddings learned using LINE's first-or second-order proximity objective function from Fit-Hi-C or O/E chromatin interaction graph, and SI analysis shows that the O/E chromatin interaction graph is good input, and the second-order proximity is a good measure to learn the global chromatin structural features. Note that SCI [12] combines two first-order and second-order loss functions by concatenating the first-order and second-order embeddings learned separately or optimization of the joint loss function to learn embeddings. Based on our evaluation, the first-order embeddings cannot capture the global compartmentalization pattern, and the second-order embeddings better represent the subcompartments. We also tried SCI's implementation of LINE for the joint optimization, however, we got the segmentation fault error, therefore, we use the learned embeddings from the optimization of the second-order objective function implemented in the original LINE [11] paper in this paper.

We also used the SI analysis to choose LINE hyperparameters. Two main hyperparameters of the LINE are sample size and embedding size. The sample size indicates the number of sampled edges during the training, and the embedding size is a dimension of the embedding space. Fig. S 2c and Fig. S 3a show that the sample size of $50M$ results in higher SI based on GM12878 and K562 Hi-C embeddings respectively, and Fig. S 2d shows that the embedding size of 8 is enough to embed the chromatin interaction graph. Furthermore, the TSA-seq signals [18] that capture the distance of genomic regions to nuclear components such as LaminA are available for the K562 cell type. We expect good structural embeddings and annotations to be highly correlated with such signals. Therefore, we used K562 Hi-C embeddings learned with different hyperparameters to identify HMM_structural domain types, and calculated the variance explained for the LaminA TSA-seq signal based on those annotations. Fig. S 3b confirms that the sample size of $50M$ and embedding size of 8 are enough to capture the structural information in Hi-C data.

Due to the similarity of our embedding approach to the SCI [12], we provide a comparison with 5 SCI compartments. The SCI constructs the inter-chromosomal Hi-C graph including all the inter-chromosomal interactions, while, we also use intra-chromosomal interactions in the graph, and remove the effect of local biases by the method mentioned in supplementary information section 2. We have shown that the dimension 8 is enough to capture the compartmentalization pattern (Fig. S 2d), while SCI learns embeddings with size 100. Note that the direct comparison between our and SCI's learned embeddings is not possible, because the direct interpretation of each embedding dimension is not possible, as the embedding method, LINE, is a shallow embedding, and no function is learned to be interpretable. Therefore, we compare HMM_structural with 5 domain types to SCI's reported 5 compartments (Fig. S 4), and show that the domain types are highly overlapped and HMM_structural is significantly better than SCI based on different evaluation approaches.

### 4 The overlap and comparison of different annotations

To compare our annotations with well-known chromatin annotations such as subcompartments, we calculated the genome-wide fold-change of each domain type from our annotation over each domain type from another annotation. Given domain types $(l_1, ..., l_m)$ from the first annotation and domain types $(l'_1, ..., l'_n)$ from the second annotation, we calculate the coverage ratio of each domain type $l$ in a genome, $p(l)$. The expected ratio of genomic bins annotated with pair of domain types $(l_i, l'_j)$ is $p(l_i) * p(l'_j)$, and we count the genomic bins annotated with pair of domain types $(l_i, l'_j)$ as $O(l_i, l'_j)$. The fold-change of a domain type $l_i$ over a domain type $l'_j$ is then calculated as $\frac{O(l_i,l'_j)}{N*p(l_i)*p(l'_j)}$.

To report one statistic for the comparison of two annotations, we calculated the similarity between annotations using Adjusted Rand Index (ARI) score [19].

### 5 Bootstrapping the evaluation metrics

#### 5.1 Bootstrapping the variance explained

We use the bootstrap method to estimate the standard errors of variance explained measures, inspired by the same process for $R^2$ measure [20]. As described in section 2.5.1, given a genome annotation $a_{1:n} \in \{1...K\}^n$ ($K$ is the number of labels) and a signal vector $s_{1:n} \in \mathbb{R}^n$, the mean value of the signal over positions with label $k$ is calculated as

$$\mu_k = \frac{\sum_{i=1}^{n} \mathbb{1}(a_i = k) s_i}{\sum_{i=1}^{n} \mathbb{1}(a_i = k)}.$$

and, the predicted signal vector is $s_i^p = \mu_{a_i}$.

The difference between the signal vector and the predicted signal vector is calculated as $d_{1:n} = s_{1:n} - s_{1:n}^p$. Then, we randomly draw a bootstrap sample of size $n$, $d_{1:n}^*$ from $d_{1:n}$ with replacement. The corresponding bootstrap sample of the signal vector is calculated as $s_{1:n}^* = s_{1:n}^p + d_{1:n}^*$. Then, the bootstrap VE is computed as

$$\text{bootstrap VE} = 1 - \frac{\sum_{i=1}^{n} (d_i^*)^2}{\sum_{i=1}^{n} (s_i^* - \overline{s*})^2}.$$

This process is repeated 20 times to report 20 bootstrap VEs that can capture the standard error of reported VEs.

#### 5.2 Bootstrapping the chromatin loop OE scores

We also use the bootstrap method to estimate the standard errors of loops OE scores (observed over expected enrichment of loops). Given a ChIA-PET dataset including $n$ loops, we randomly draw a bootstrap sample of size $n$ from loops with replacement, and calculate the corresponding OE score as mentioned in the section 2.5.4. We repeat this process 20 times to report 20 bootstrap OE scores.

### 6 Existing SAGA methods

#### 6.1 Existing SAGA methods for modeling 1D signals

##### GMM

A Gaussian Mixture model is an unsupervised soft clustering method that assumes data are drawn independently from a mixture of Gaussian distributions. More formally, given a data $X = \{X_1...X_N\}$ of size $N$ and assuming that data is comprised of $K$ clusters, a latent variable $Z_i \in \{1...K\}$ is defined for each data point $X_i \in X$, where $Z_i = k$ if data point $i$ is from cluster $k$. Each cluster $k \in \{1...K\}$ is characterized by $\mu_k$ and $\Sigma_k$, the mean and covariance of its corresponding Gaussian distribution, and $\pi_k$, which is a mixing probability of a cluster. Then, the probability that $X_i$ is from cluster $k$ is

$$p(X_i, Z_i = k) = p(X_i|Z_i = k)p(Z_i = k) = \mathcal{N}(X_i|\mu_k, \Sigma_k) * \pi_k,$$

where $p(X_i|Z_i = k)$ is an emission probability of data point $i$ from cluster $k$, and $p(Z_i = k)$ is a constant representing the mixing probability of cluster $k$. The joint probability of all observed variables $X$ and latent variables $Z$ is defined as

$$p(X, Z) = \prod_{i=1}^{N} P(X_i|Z_i)P(Z_i).$$

Given a model learned parameters $\mu_k$, $\Sigma_k$, and $\pi_k$ for each cluster $k$, the posterior distribution over $K$ clusters for each data point is calculated during inference to infer the most probable cluster.

**GMM-HMM**

Hidden Markov model adds the dependency between neighbor latent variables to GMM's naive probabilistic model. So, the probability that $X_i$ is from cluster $k$ is defined as $p(X_i, Z_i = k) = p(X_i|Z_i = k)p(Z_i = k|Z_{i-1})$, and the joint probability of all observed variables $X$ and latent variables $Z$ would be:

$$p(X, Z) = \pi_{Z_1} p(X_1|Z_1) \prod_{i=2}^{N} P(X_i|Z_i)P(Z_i|Z_{i-1}),$$

where $P(Z_i = z_i|Z_{i-1} = z_{i-1})$ is captured in a transition probability distribution $A(z_i|z_{i-1})$.

**Segway**

The Segway model [2] incorporates additional types of latent variables which are specifically designed for modelling genomic signals. The main feature of Segway is modeling the distribution of segments length by putting a prior on segments length distribution. This is done by determining a prior on the range of segments lengths that is modeled by the transition model, and assigning weight to the transition model, such that higher weights give more power to the transition model than the emission model during learning and inference.

#### 6.2 Existing integrative SAGA methods for jointly modeling 1D signals and spatial dependencies

**SPIN (HMRF)**

A Markov random field adds higher-order dependencies between latent variables to the naive probabilistic model in GMM. The dependency between latent variables is captured in a Markov random field represented as a graph $G = (V, E)$, where $V = \{V_1 ... V_N\}$ are corresponding to latent variables $Z = \{Z_1 ... Z_N\}$ and $(i, j) \in E$ if there is a dependency between $Z_i$ and $Z_j$ (SPIN [9] uses the neighboring genomic bins in the linear genome and 3D space to construct the random field). $p(X_i|Z_i)$ is similar to the emission probability distribution in GMM, and $p(Z_i, Z_j)$ is a transition probability distribution to model the dependency between all neighboring genomic bins. Then, the joint probability of all observed variables $X$ and latent variables $Z$ is defined as

$$p(X, Z) = \prod_{i=1}^{N} P(X_i|Z_i) \prod_{(i,j)\in E} p(Z_i, Z_j).$$

**Segway-GBR**

GMM-HMM and HMRF incorporate the dependency between posterior probability distributions of neighbor latent variables in the definition of the joint probability distribution $P(X, Z)$. A GMM-HMM cannot model higher-order dependencies, and HMRF inference is done by approximate algorithms which are not guaranteed to converge, specifically when there is not a high agreement between observed variable and the MRF structure. Therefore, graph-based posterior regularization (GBR) [8] was proposed to incorporate the higher-order dependency information by augmenting a probabilistic model like Segway given a regularization graph such that nearby variables in a regularization graph have similar posterior distributions. This is done by adding a regularizer term to the base probabilistic model's loss. The regularizer can be added either during training to learn both model parameters and posterior distributions with respect to the regularizer graph, or during the inference to infer the posterior distributions given the original model parameters with respect to the regularizer graph. We train a regular Segway, and use GBR during inference similar to the Segway-GBR [21]. While the learning and inference of probabilistic models is based on maximizing the objective which is equal to the likelihood of observed data $\mathcal{L}(\theta) = log\ p_\theta(X)$, GBR adds a regularizer term to the objective to encourage the similar posterior distributions for neighboring latent variables:

$$\begin{aligned}\text{maximize}_{\theta,q}\ \ \mathcal{J}(\theta, q) = \mathcal{L}(\theta) - D(q(Z) \parallel p_\theta(Z|X)) \\ - \lambda_G \sum_{(u,v)\in E_R} w(u, v) D(q_u^M(Z_u) \parallel q_v^M(Z_v)),\end{aligned}$$

where $q(Z)$ is an auxiliary joint distribution over latent variables $Z$, $q_u^M(Z_u)$ is a marginal distribution over a latent variable $Z_u$, and $D$ is the Kullback-Leibler divergence. The second term encourages the auxiliary joint distribution and inferred joint distribution over latent variables (given observed variables) to be similar, and the third term encourages the marginal distribution of neighbor latent variables in a regularization graph to be similar. $E_R$ is the set of edges of the regularization graph, $w(u, v)$ is the weight of edge between latent variables $Z_u$ and $Z_v$, and $\lambda_G$ controls the strength of regularization.

### 7 SPIN result analysis

We found that SPIN is not effective when using functional genomic signals together with Hi-C significant interactions as input, and usually assigns the whole genome to one or several labels while the other labels cover no positions. However, it performs well when input with spatial genomic signals in the original paper, LaminB and SON TSA-seq and LaminB and Nucleolus DamID.

In contrast to spatial signals, functional signals, fluctuate rapidly. Thus, the naive clustering of structural signals with GMM without spatial constraint results in stable annotation, while the annotation by applying GMM on functional signals is very fluctuating (Fig. S 9a). The performance of the loopy belief propagation-based model, SPIN (HMRF), is highly dependent on the stability of the neighborhood information encoded in its Markov random field (MRF). For example, we found that HMRF with functional signals and chain edges as an input results in a solution with only a few labels, and dominated with one label (Fig. S 9b), since the sequential pattern of functional states is very noisy (Fig. S 9a).

We found that adding Hi-C significant interactions to the MRF as done by SPIN can reduce such noise by increasing the degree of nodes in the MRF, and capturing the stable pattern in the nodes' neighborhood information. For example, adding a sufficient number of significant interactions result in MRF that separates active and inactive nodes. However, the performance is highly dependent on the choice of a number of significant interactions, as the small numbers of Hi-C interactions are noisy because of the low degree of nodes, and the large numbers are noisy because they are not necessarily significant.

We showed this behavior by an experiment on the chromosome 16 and inputting functional signals with Hi-C significant interactions to SPIN. We did it with a different number of Hi-C significant interactions, and two types of measurement to choose significant interactions, lower Fit-Hi-C p-values and higher normalized Hi-C counts (observed/expected (O/E) values), in 3 different resolutions. The goodness of annotations is shown as a similarity with well-known annotations, subcompartments [5], A/B compartments, and GMM_functional domain types (Fig. S 9c).

We can see that a small and large number of significant interactions result in invalid annotations with a near-zero agreement with well-known annotations. We noticed that valid annotations from SPIN are also two-state annotations that separate active and inactive regions but do not distinguish further subtypes. This can be seen as a high agreement with A/B compartments. The number of proper significant interactions is dependent on the resolution, such that higher-resolution settings with more nodes also require more edges to capture the structural organization pattern. The performance also depends on the type of significant interactions, for example, Fit-Hi-C significant interactions in higher resolutions, 10 kb, do not include very many long-range interactions that discriminate A/B compartments, therefore, increasing the number of significant interactions does not improve the performance. Lastly, we visualized the coverage of domain types from invalid and valid SPIN annotations (Fig. S 9b), and we can see that the invalid solution is almost covered with one state, and does not explain any type of cellular activity and chromatin features, and the valid solution is a two-states annotation that separate active and inactive chromatin types.

#### 8 Comparison of combinatorial and paired domain types

An alternative strategy for integrative domain annotation is to use the pair of HMM_functional and HMM_structural domain types as combined domain types, resulting in $6 * 6 = 36$ labels (also known as late integration in ML area). We produced HMM_combined domain types with $k = 36$ to compare to paired labels, named HMM_joint. The HMM_combined_k36 annotation is significantly better than HMM_joint_k36 in 3/4 evaluation tasks, and it is comparable to HMM_joint_k36 in terms of proportion of variance explained for replication timing signals (Fig. S 17b). Furthermore, the purpose of integrative genome annotation in different scales is to summarize a variety of genomic data into a limited number of features or labels to be more interpretable. However, lots of paired labels are rare and only 18 pairs have more than 1% coverage (Fig. S 17a). Therefore, the early integration approach to summarize all data sets into a few domain types is more easier to interpret than pairing different annotations.

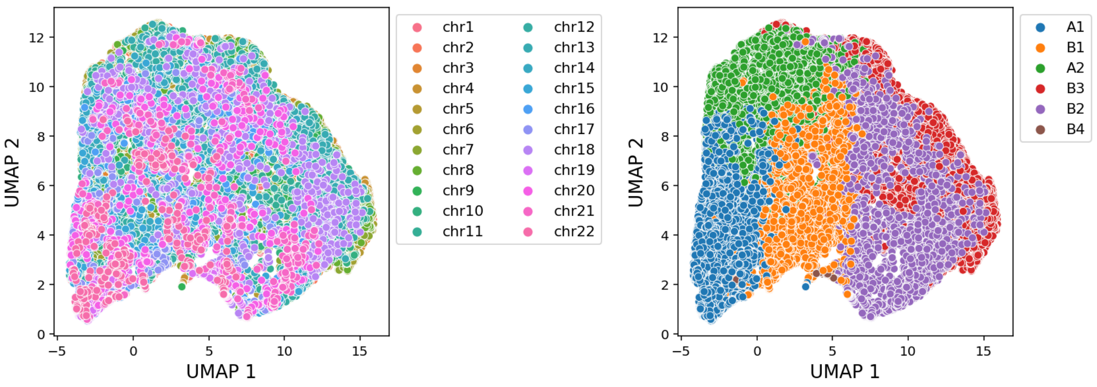


(a)

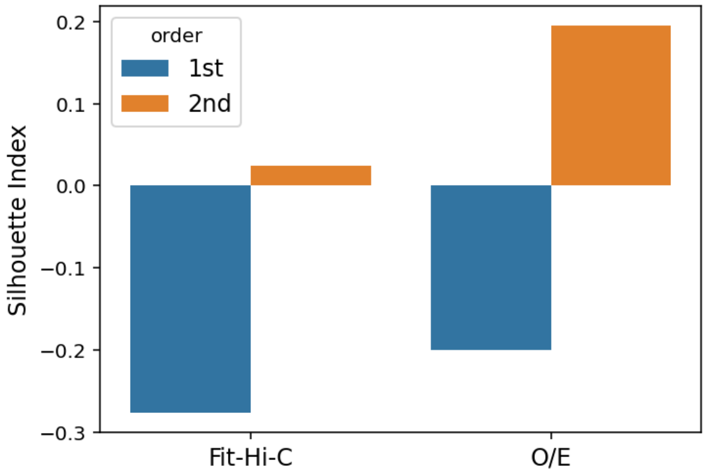


(b)

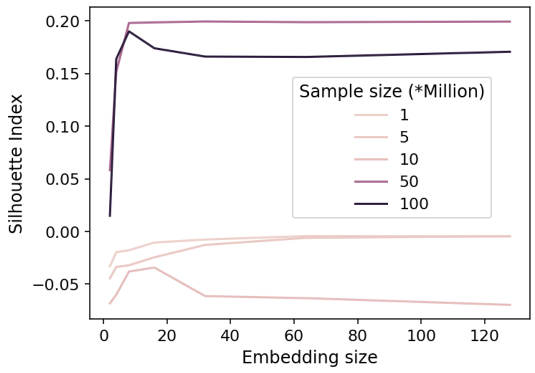


(c)

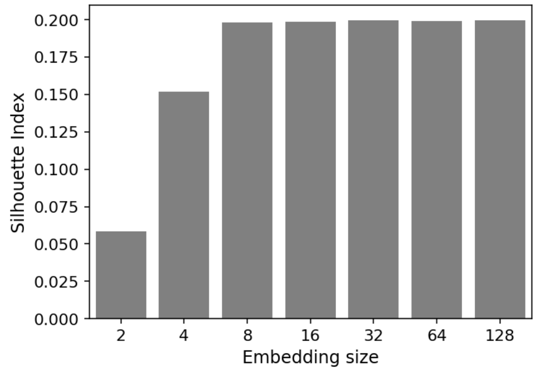


(d)

Fig. S2: (a) The scatter plots of LINE embeddings (2nd order) UMAPs for the O/E chromatin interaction graph colored by chromosome numbers (left) and subcompartment labels (right). (b) Silhouette Index of learned embeddings from Fit-Hi-C and O/E chromatin interaction graphs by 1st and 2nd orders objectives according to subcompartment labels. (c) Silhouette Index of 2nd-order learned embeddings from O/E chromatin interaction graph according to subcompartment labels with respect to two LINE hyperparameters, sample size (number of sampled edges during training), and embedding size. (d) Silhouette Index of 2nd-order learned embeddings from O/E chromatin interaction graph according to subcompartment labels for different embedding sizes (sample size = 50 Million). This plot is for a cell type, GM12878.

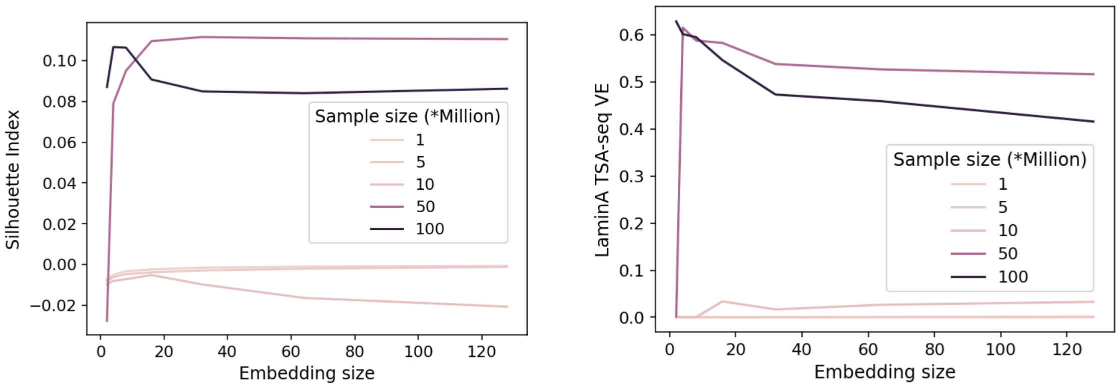


(a) (b)

Fig. S3: (a) Silhouette Index of 2nd-order learned embeddings from O/E chromatin interaction graph according to subcompartment labels with respect to two LINE hyperparameters, sample size (number of sampled edges during training), and embedding size. (b) The proportion of variance explained of LaminA TSA-seq signal according to HMM_structural domain types identified by inputting learned embeddings from O/E interaction graph to HMM with respect to two LINE hyperparameters. This plot is for a cell type, K562.

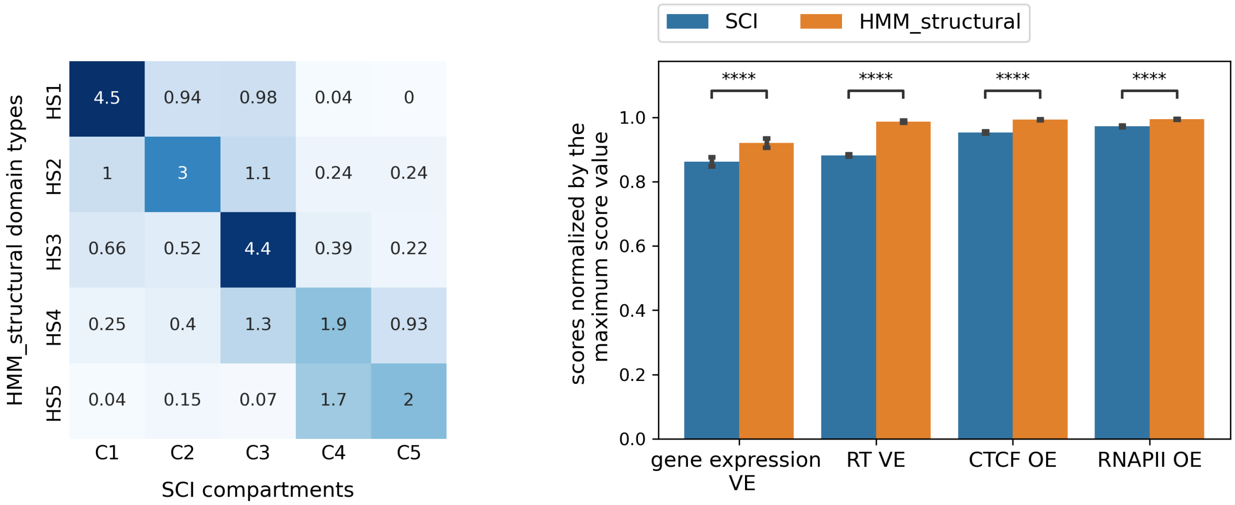


Fig. S4: The comparison between HMM_structural ($k = 5$) annotation and SCI [12] annotation through their fold-change overlap (left), and based on the proportion of variance explained for gene expression (gene expression VE), the average proportion of variance explained for replication timing 6 phases signals (RT VE), and agreement with CTCF and RNAPII ChIA-PET loops (CTCF OE, RNAPII OE) (right).

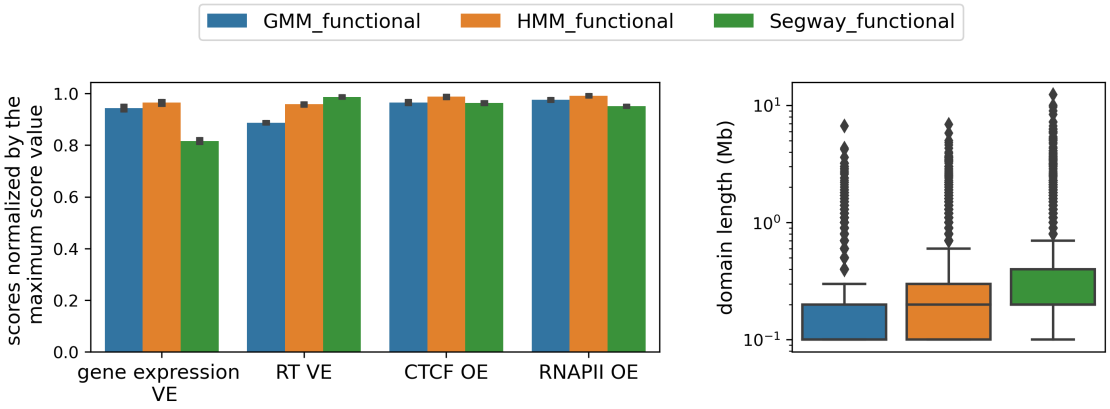


(a)

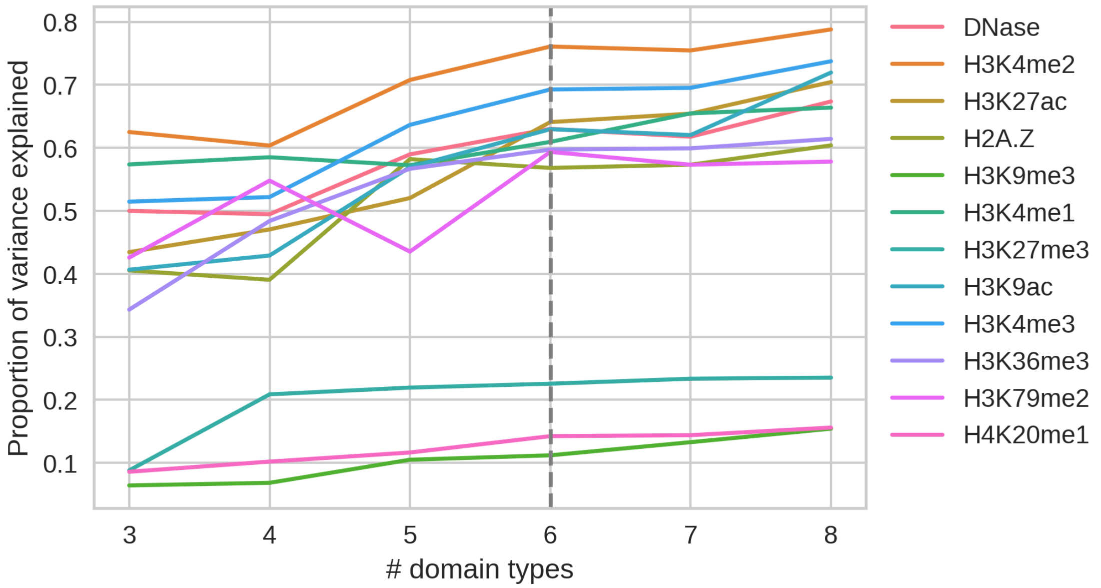


(b)

Fig. S5: (a) The comparison between GMM_functional, HMM_functional, and Segway_functional annotations based on the proportion of variance explained for gene expression (gene expression VE), the average proportion of variance explained for replication timing 6 phases signals (RT VE), agreement with CTCF and RNAPII ChIA-PET loops (CTCF OE, RNAPII OE), and the distribution of domain lengths. (b) The proportion of variance explained for input signals according to HMM_functional annotations with different numbers of domain types. This plot is for a cell type, GM12878.

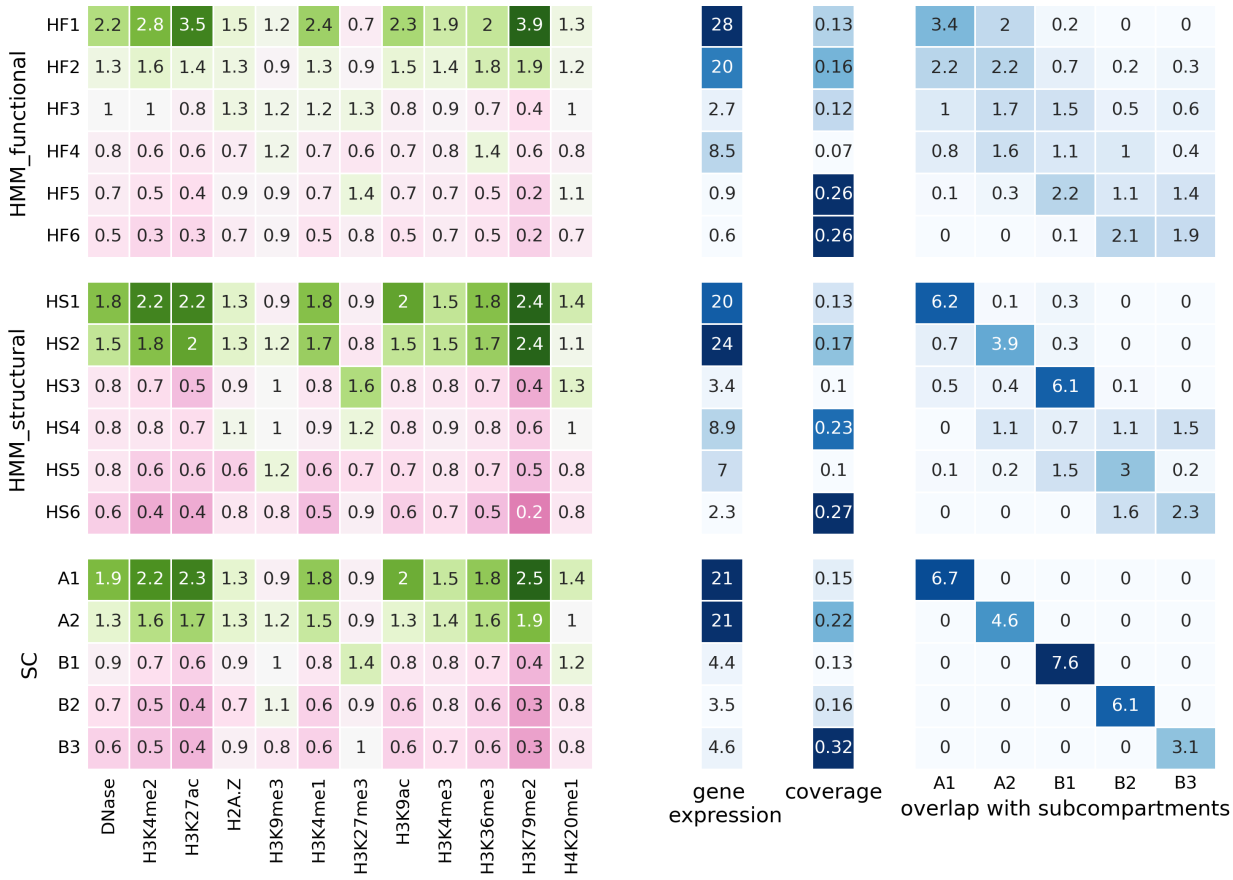


Fig. S6: The fold enrichment of genomic signals (column 1), average of transcription level of genes (column 2), coverage (column 3), and fold-change of subcompartments (column 4) of each domain type from HMM_functional, HMM_structural, and subcompartments annotations. This plot is for a cell type, GM12878.

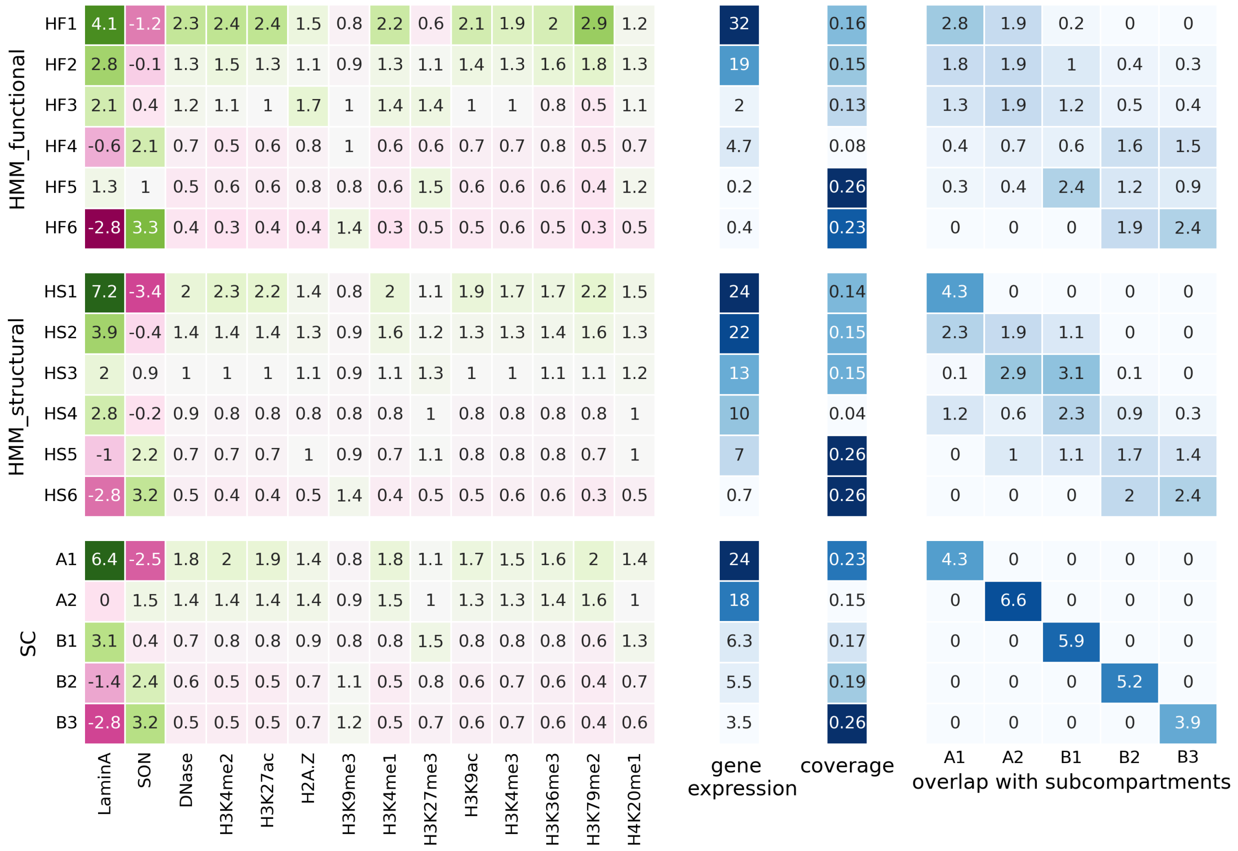


Fig. S7: The fold enrichment of genomic signals (column 1), average of transcription level of genes (column 2), coverage (column 3), and fold-change of subcompartments (column 4) of each domain type from HMM_functional, HMM_structural, and subcompartments annotations. This plot is for a cell type, K562.

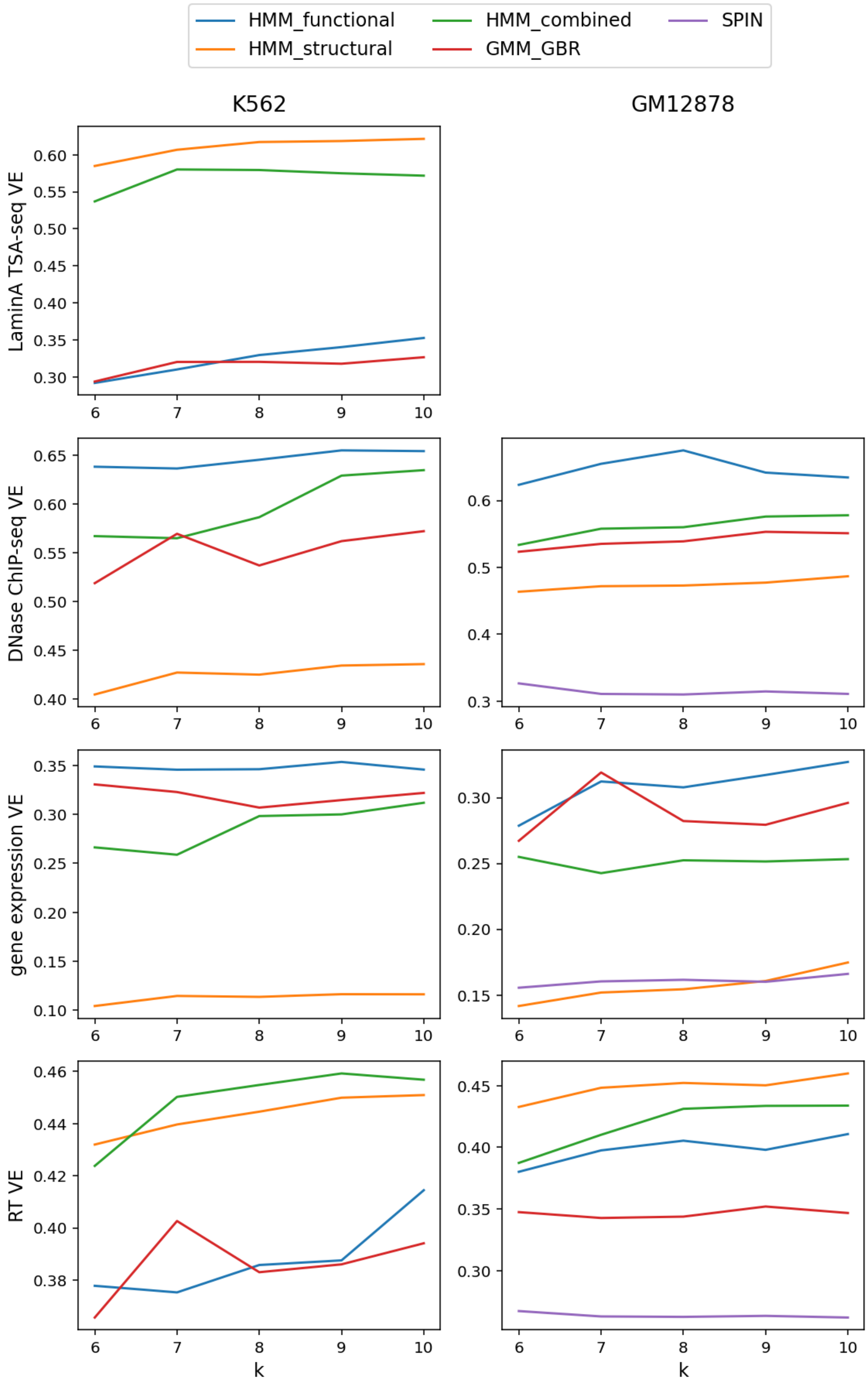


Fig. S8: The proportion of variance explained for LaminA TSA-seq, DNase ChIP-seq, gene expression, and replication timing signals given HMM_functional, HMM_structural, and HMM_combined annotations with a number of domain types (k) from 6 to 10.

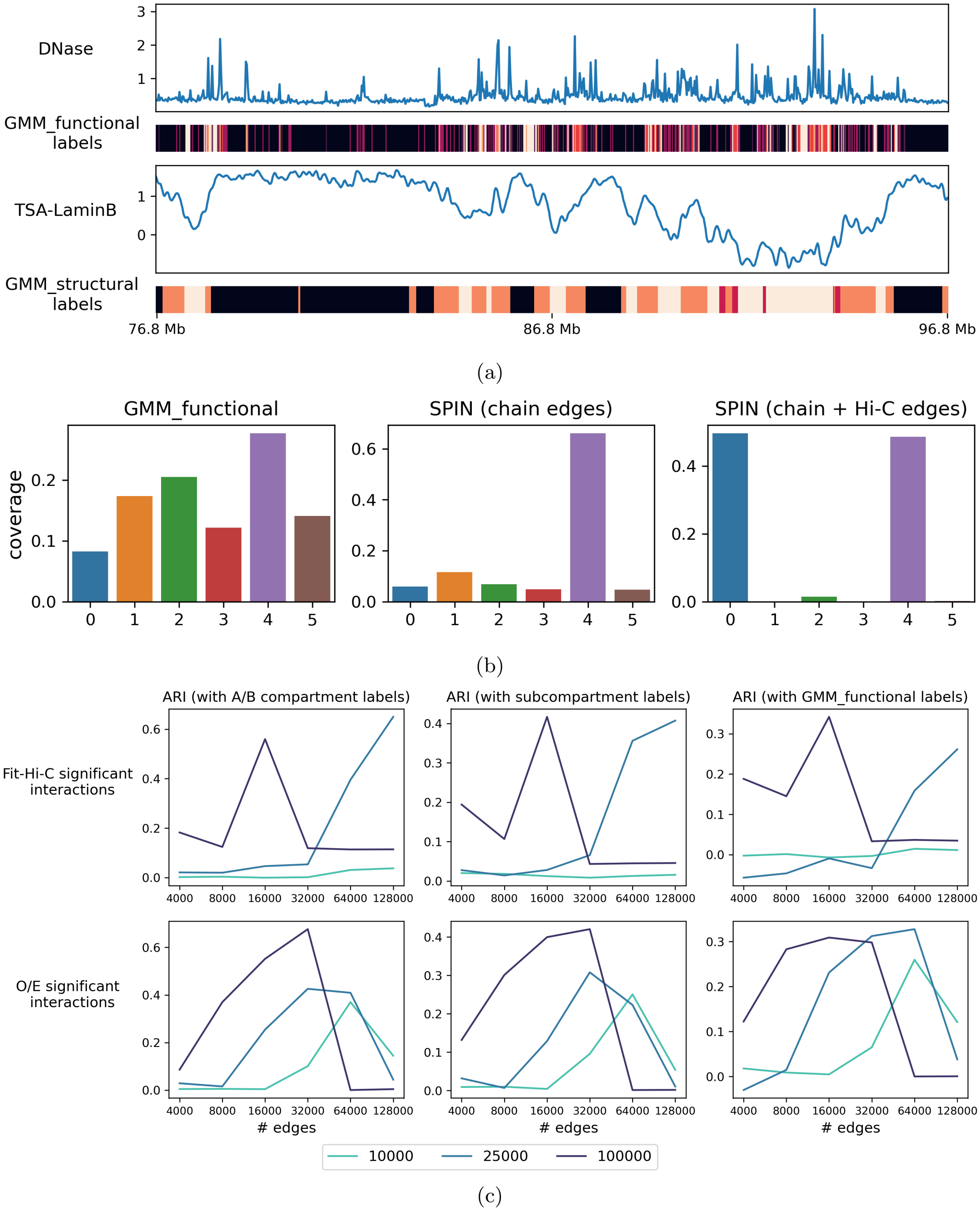


Fig. S9: (a) Comparison of a functional genomic signal, DNase, and a structural genomic signal, TSA-LaminB, and annotations based on functional and structural genomic signals (GMM_functional and GMM_structural labels respectively). (b) Coverage of the GMM_functional, SPIN with chain and chain+Hi-C edges domain types. (c) The similarity of SPIN annotation (given our data) with A/B compartments, subcompartments, and GMM_functional annotations based on the number and type of input significant interactions in different resolutions.

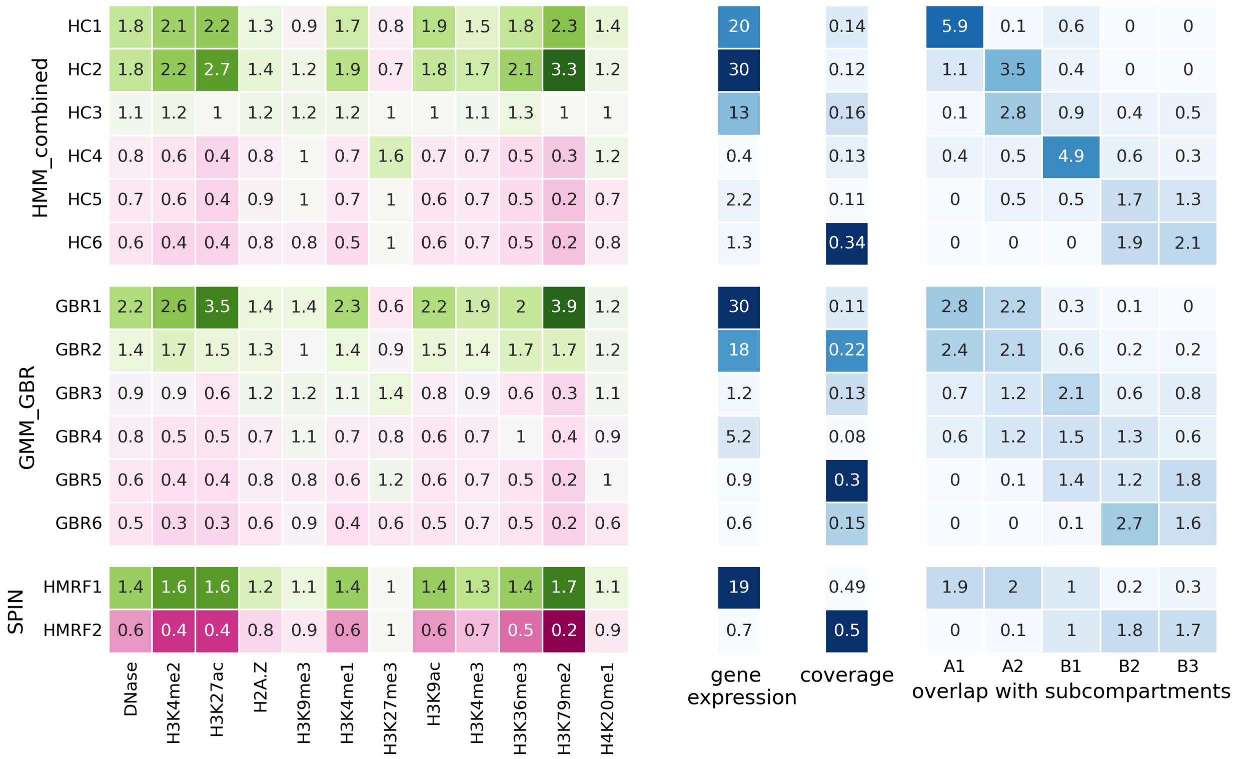


Fig. S10: The fold enrichment of genomic signals (column 1), an average of transcription level of genes (column 2), the density of genes (column 3), coverage (column 3), and fold-change of subcompartments of each domain type of HMM_combined, GMM_GBR, and SPIN annotations. This plot is for a cell type, GM12878.

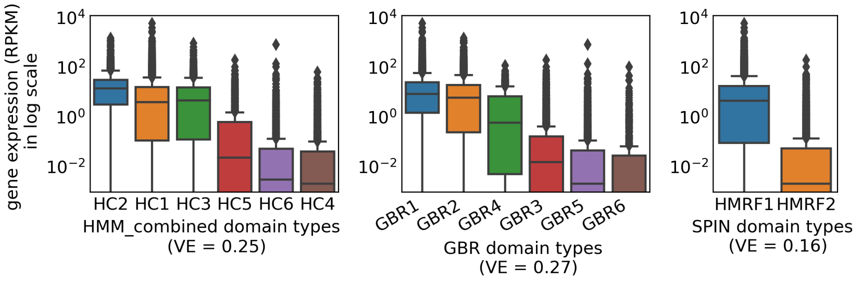


(a)

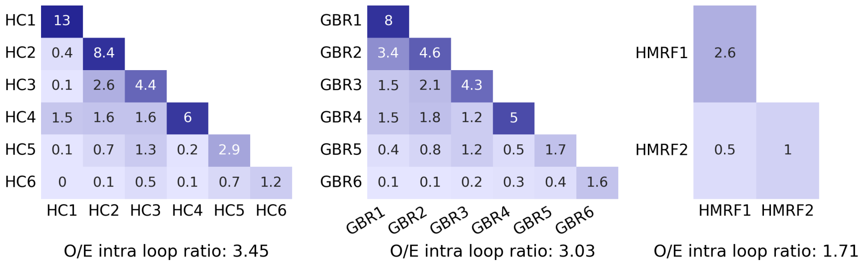


(b)

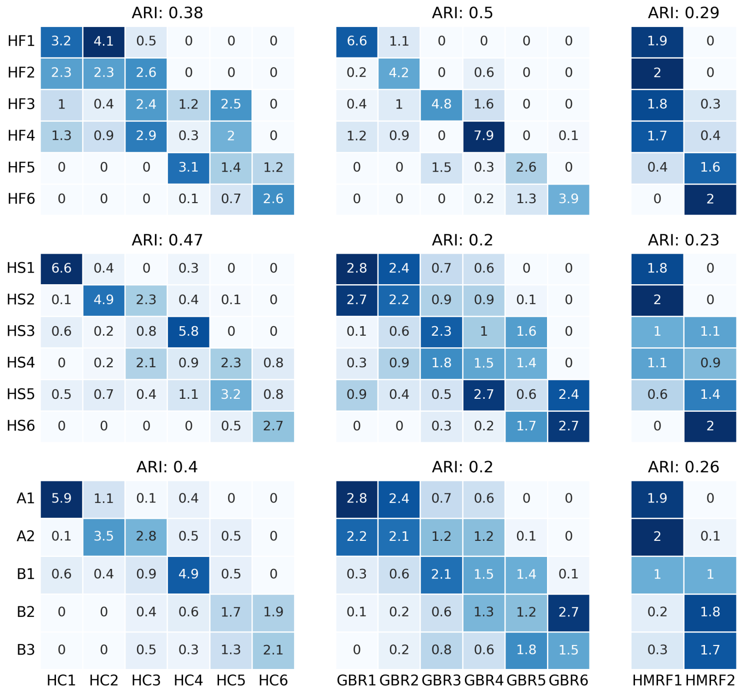


(c)

Fig. S11: (a) The distribution of gene expression values in RPKM for each domain type of HMM_combined, GMM_GBR, and SPIN annotations. 'VE' in titles means the variance explained for gene expression given a domain annotation. (b) The observed/expected CTCF ChIA-PET loops between each pair of labels for different annotations. (c) The overlap of each of HMM_combined, GMM_GBR, and SPIN domain types with HMM_functional, HMM_structural, and subcompartments domain types and similarity (adjusted rand index (ARI)) between the annotations. This plot is for a cell type, GM12878.

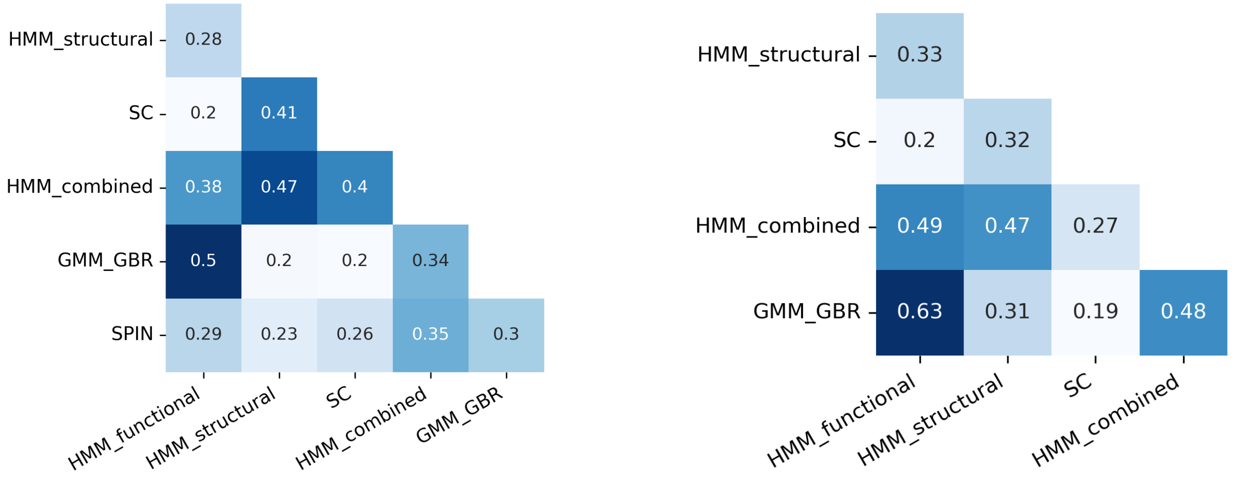


(a) (b)

Fig. S12: The similarity (adjusted rand index (ARI)) between all pairs of annotations for (a) GM12878 and (b) K562 cell types.

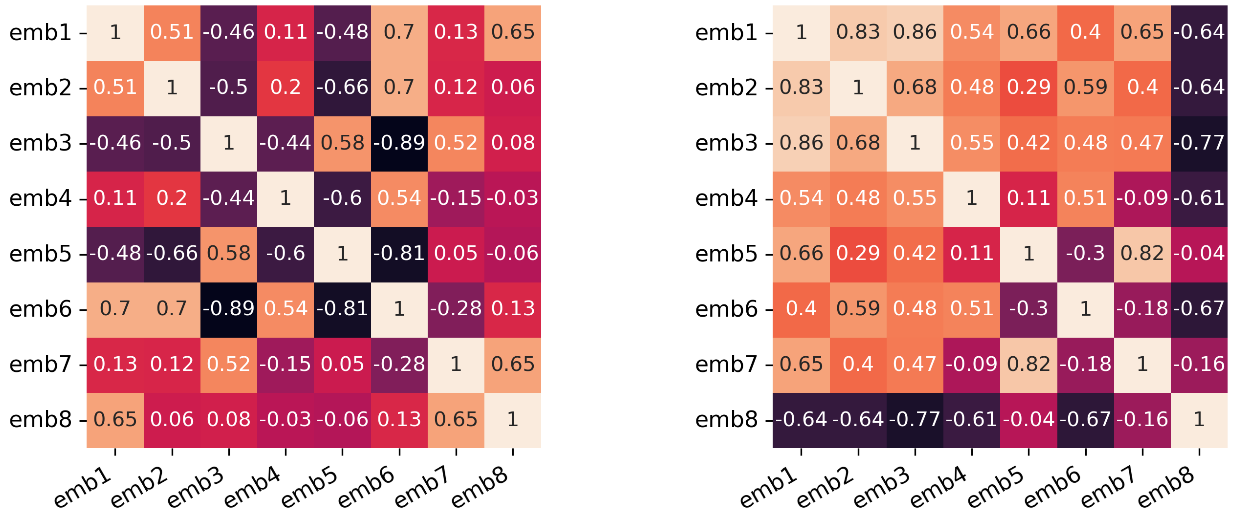


(a) (b)

Fig. S13: The correlation between dimensions of learned structural features for (a) GM12878 and (b) K562 cell types.

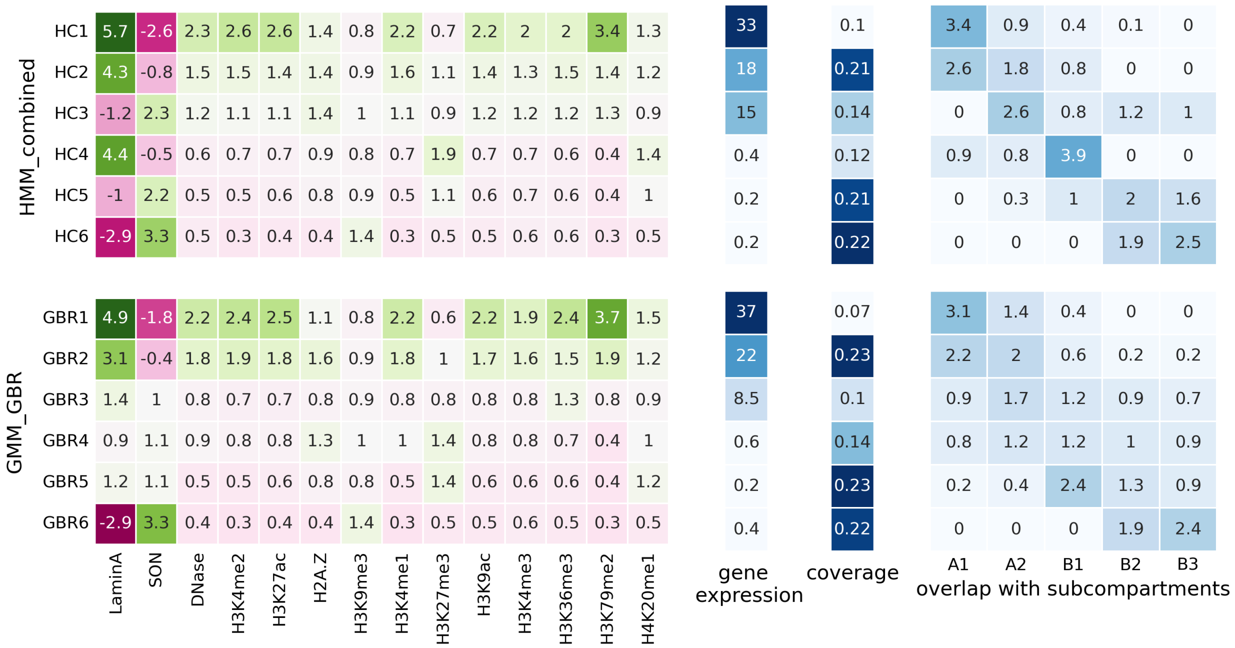


(a)

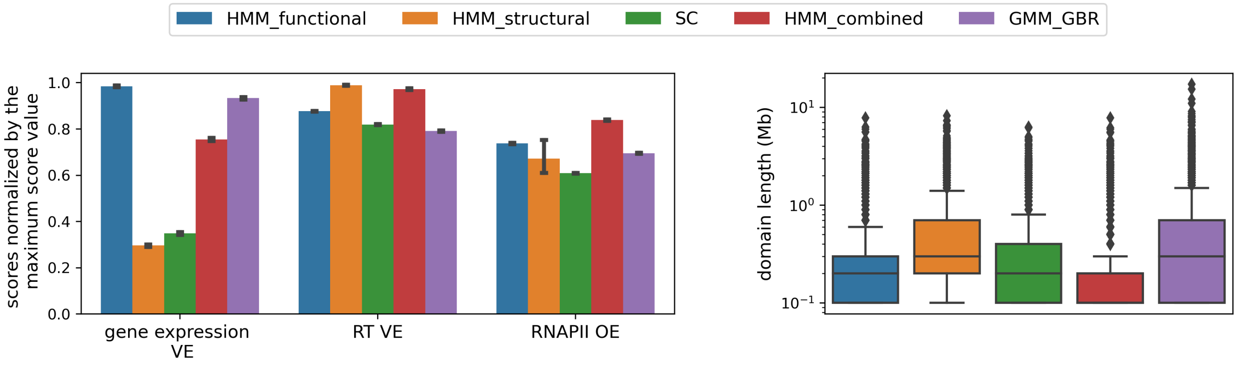


(b)

Fig. S14: (a) The fold enrichment of genomic signals (column 1), an average of transcription level of genes (column 2), the density of genes (column 3), coverage (column 3), and fold-change of subcompartments of each domain type of HMM_combined and GMM_GBR. (b) Comparison of HMM_functional, HMM_structural, subcompartments, HMM_combined, and GMM_GBR annotations based on the proportion of variance explained for gene expression (gene expression VE), the average proportion of variance explained for replication timing 6 phases signals (RT VE), agreement with RNAPII ChIA-PET loops (RNAPII OE), and the distribution of domain lengths. This plot is for a cell type, K562.

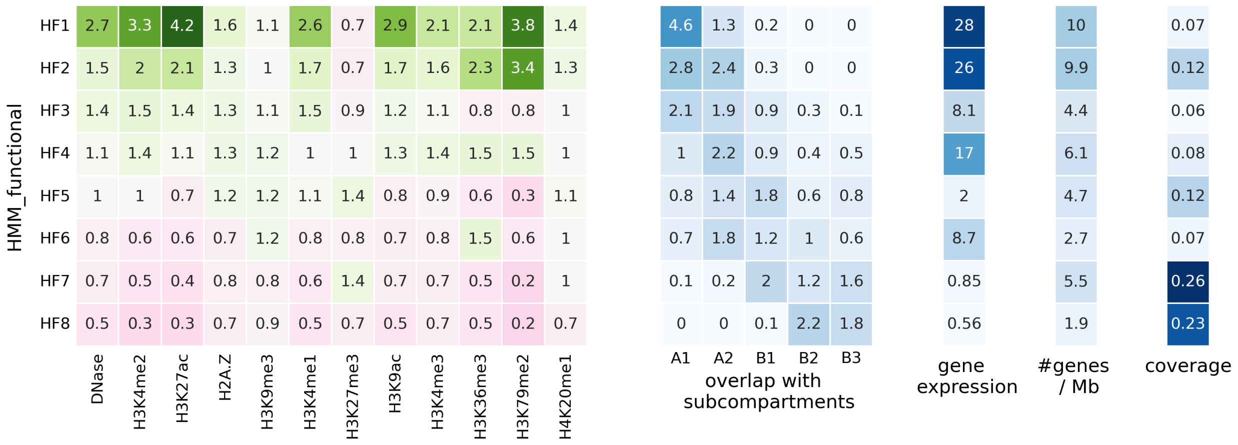


(a)

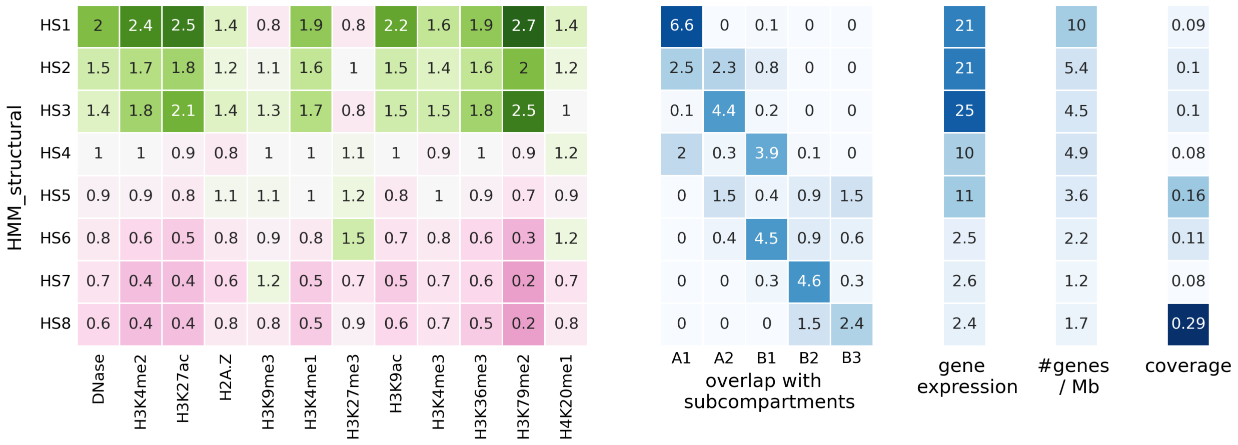


(b)

Fig. S15: The fold enrichment of genomic signals (column 1) and subcompartments (column 2), an average of transcription level of genes (column 3), the density of genes (column 4) and coverage (column 5) of each of 8 domain types from (a) HMM_functional and (b) HMM_structural annotations. This plot is for a cell type, GM12878.

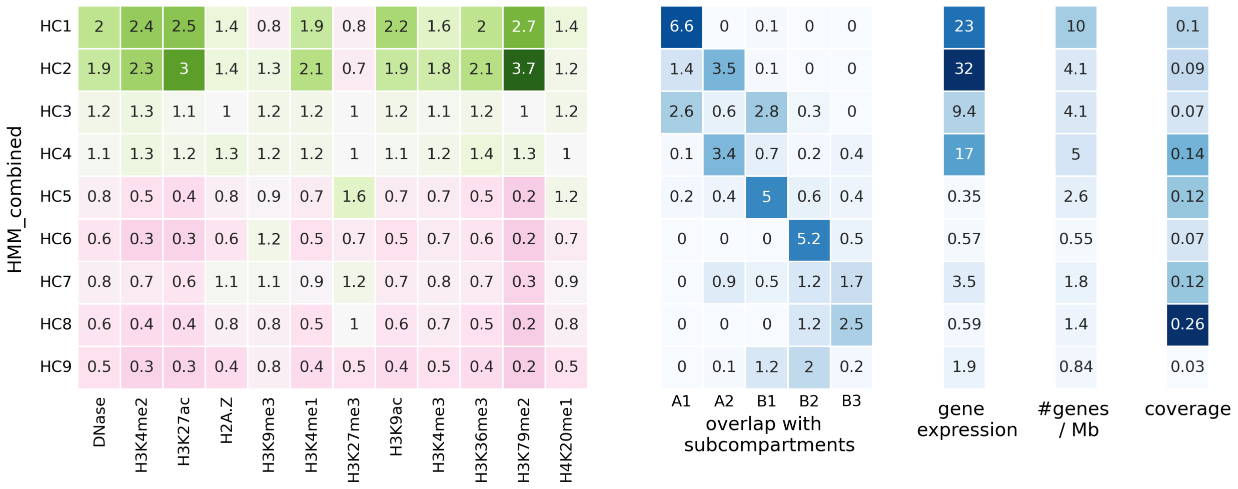


(a)

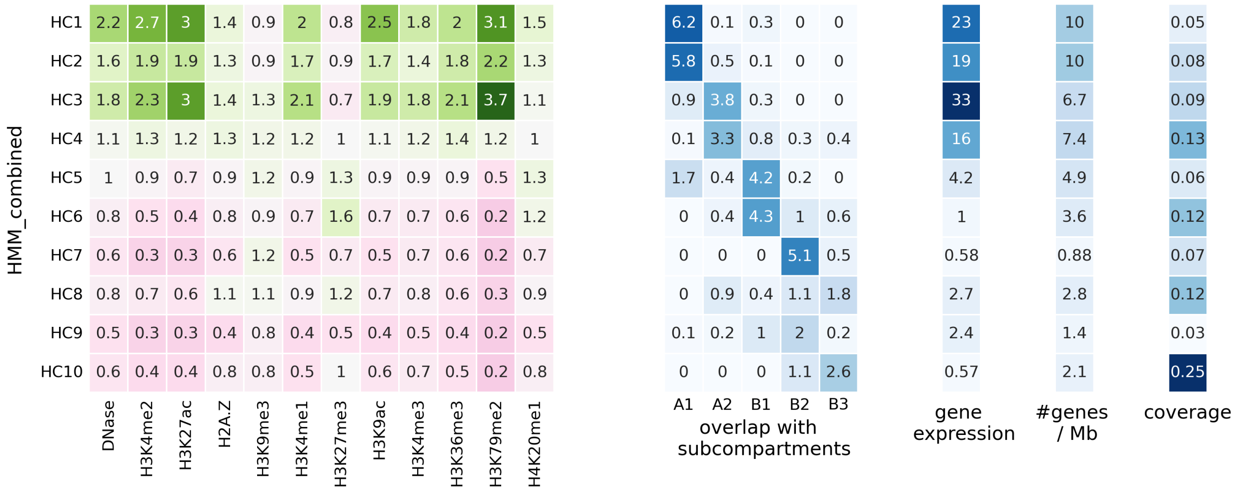


(b)

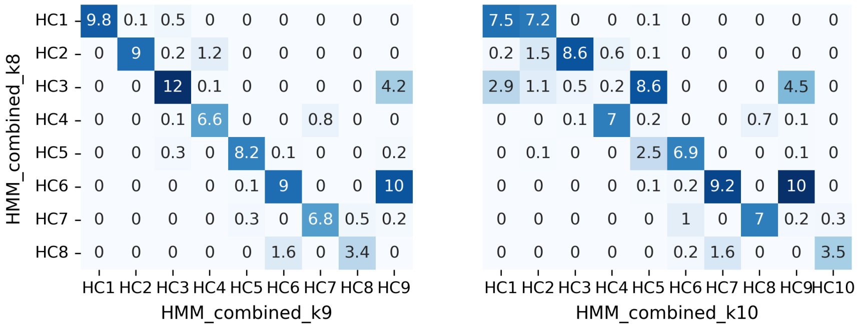


(c)

Fig. S16: The fold enrichment of genomic signals (column 1) and subcompartments (column 2), an average of transcription level of genes (column 3), the density of genes (column 4) and coverage (column 5) of each domain type from (a) HMM_combined_k9 and (b) HMM_combined_k10 annotations. (c) The fold-change overlap between domain types of HMM_combined_k8 annotation with domain types of HMM_combined_k9 (left) and HMM_combined_k10 (right) annotations. This plot is for a cell type, GM12878.

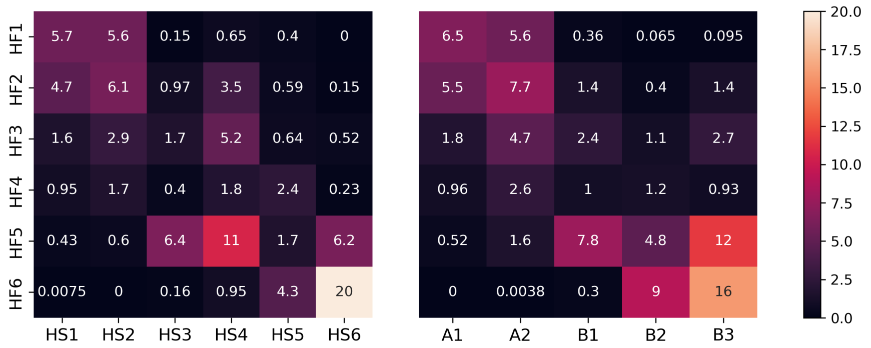


(a)

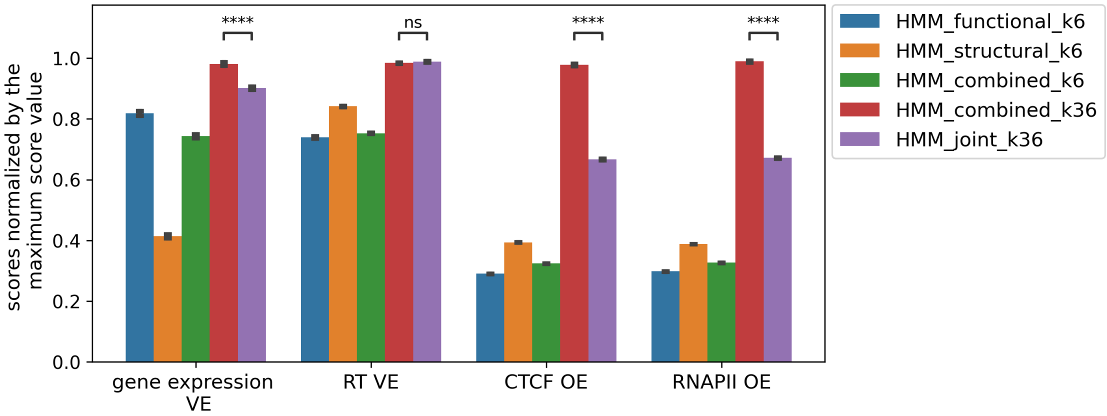


(b)

Fig. S17: (a) The percentage of bins annotated with pairs of HMM_functional and HMM_structural domain types (left), and pairs of HMM_functional and subcompartment domain types (right) (b) The comparison of HMM_combined with $k = 36$ and HMM_joint which is the annotation based on paired annotations from HMM_functional and HMM_structural.

[7] Kyle Xiong and Jian Ma. Revealing hi-c subcompartments by imputing inter-chromosomal chromatin interactions. *Nature communications*, 10, 2019.

[16] Leland McInnes, John Healy, and James Melville. Umap: Uniform manifold approximation and projection for dimension reduction. *arXiv preprint arXiv:1802.03426*, 2018.
